# The *S. cerevisiae* m^6^A-reader Pho92 impacts meiotic recombination by controlling key methylated transcripts

**DOI:** 10.1101/2022.03.21.485107

**Authors:** Jérémy Scutenaire, Damien Plassard, Mélody Matelot, Tommaso Villa, Julie Zumsteg, Domenico Libri, Bertrand Séraphin

## Abstract

N^6^-methyladenosine (m^6^A), the most abundant internal modification of eukaryotic mRNAs, participates in the post-transcriptional control of gene expression. In *Saccharomyces cerevisiae*, m^6^A is only found during meiosis. Although the deletion of the m^6^A- methyltransferase Ime4 impairs this process, the molecular impact of m^6^A on gene expression remains ill defined. Here we investigated the function of the budding yeast m^6^A reader Pho92. We found that Pho92 is specifically expressed during meiosis and impacts meiotic progression. We used high-throughput RNA sequencing and mapping of Pho92-binding sites following UV-crosslinking to show that Pho92 is recruited to specific mRNAs in an m^6^A-dependent manner during the meiotic prophase, preceding their down-regulation. Strikingly, point mutations altering m^6^A sites in mRNAs targeted by Pho92 are sufficient to delay their down-regulation and, in one case, to impact meiotic progression. Altogether, our results indicate that Pho92 facilitate the meiotic progression by accelerating the down-regulation of timely-regulated mRNAs during meiotic recombination.

## INTRODUCTION

A prerequisite of life is the fast and efficient ability of organisms to adjust their cellular content in response to internal and external cues. This most often entails changes in gene expression that are achieved at multiple levels including transcription, RNA processing, translation and RNA decay. Recently, epitranscriptomics, that involves the regulated post-transcriptional modifications of RNA bases, emerged as a novel fine-tuning layer of gene expression (Zhao et al., 2016; Seo and Kleiner, 2021). N^6^-adenosine methylation (m^6^A) is the most prevalent internal modification of eukaryotic mRNAs and participates in multiple fundamental processes such as development, cellular differentiation and gametogenesis (Hsu et al., 2017a; Klungland et al., 2017). It also emerged as a recent therapeutical target in human neurological diseases and cancer metabolism (Chen et al., 2019; Du et al., 2019; He and He, 2021). m^6^A is found on a subset of RRACH (R: A/G, H: A/C/U) sites in a wide array of organisms, or is associated with 2’methyl-ribose as part of the vertebrate mRNA cap structure (Mauer et al., 2017). The former sites are enriched in the vicinity of the stop codon in coding sequences and 3’UTR in mammals (Dominissini et al., 2012; Meyer et al., 2012), plants (Bodi et al., 2012), yeast (Schwartz et al., 2013) and flies (Lence et al., 2016). m^6^A is added cotranscriptionally on polymerase II transcribed RNAs by a multi-component “writer” complex comprising METTL3-METTL14 heterodimer in metazoans and plants (Zhong et al., 2008; Slobodin et al., 2017; Zaccara et al., 2019), of which METTL3 (also called MT-A70) is the active component (Śledź and Jinek, 2016; Wang et al., 2016a, 2016b). In the yeast *S. cerevisiae*, the METTL3 orthologue is encoded by Ime4 (Inducer of meiosis 4) and associates with 2 non-catalytic factors: Mum2 (Muddled meiosis 2, orthologous to mammalian WTAP), and Slz1 (Sporulation leucine zipper 1) (Agarwala et al., 2012; Clancy et al., 2002).

Unlike animals and plants, m^6^A has been detected in *S. cerevisiae* transcripts only during meiosis (Clancy et al., 2002; Agarwala et al., 2012; Schwartz et al., 2013). Yeast meiosis is part of the sporulation program induced in diploid cells by starvation for nitrogen in the presence of a non-fermentable carbon source that results in the production of 4 haploid spores (reviewed in Piekarska et al., 2010; Neiman, 2011). This process is controlled by the master regulatory transcription factor *IME1* (Inducer of meiosis 1) which activates a first transcriptional wave of genes sharing a common regulatory URS1 (Upstream repression sequence I) site in their promoters. These “early” genes are required for the successive events of DNA replication, programmed formation of DNA double-strand breaks (DSBs), assembly of synaptonemal complex structures that connect homologous chromosomes, and initiation of meiotic recombination (Figure 1A). A second major meiotic phase involves the transcriptional factor Ndt80 (Non-DiTyrosine) to activate the expression of “middle genes” needed to complete recombination, trigger the two meiotic nuclear divisions (meiosis I and II) and initiate spore formation. Most of the “middle” genes contain the MSE (Middle sporulation element) motif in their promoters acting as Ndt80-binding site. In non-meiotic conditions, sporulation genes are transcriptionally repressed by Ume6 (Unscheduled meiotic gene expression 6) and Sum1 (Suppressor of Mar1-1) which bind to URS1 and MSE sites, respectively. Beyond transcriptional control, entry and progression into sporulation requires multiple coordinated regulatory pathways including RNA modifications (Schwartz et al., 2013), translational regulation (Berchowitz et al., 2013; Jin and Neiman, 2016), precise post-translational modifications and fine-tuning of protein levels (Eisenberg et al., 2018; Bhagwat et al., 2021; Shi et al., 2021).

**Figure 1.**
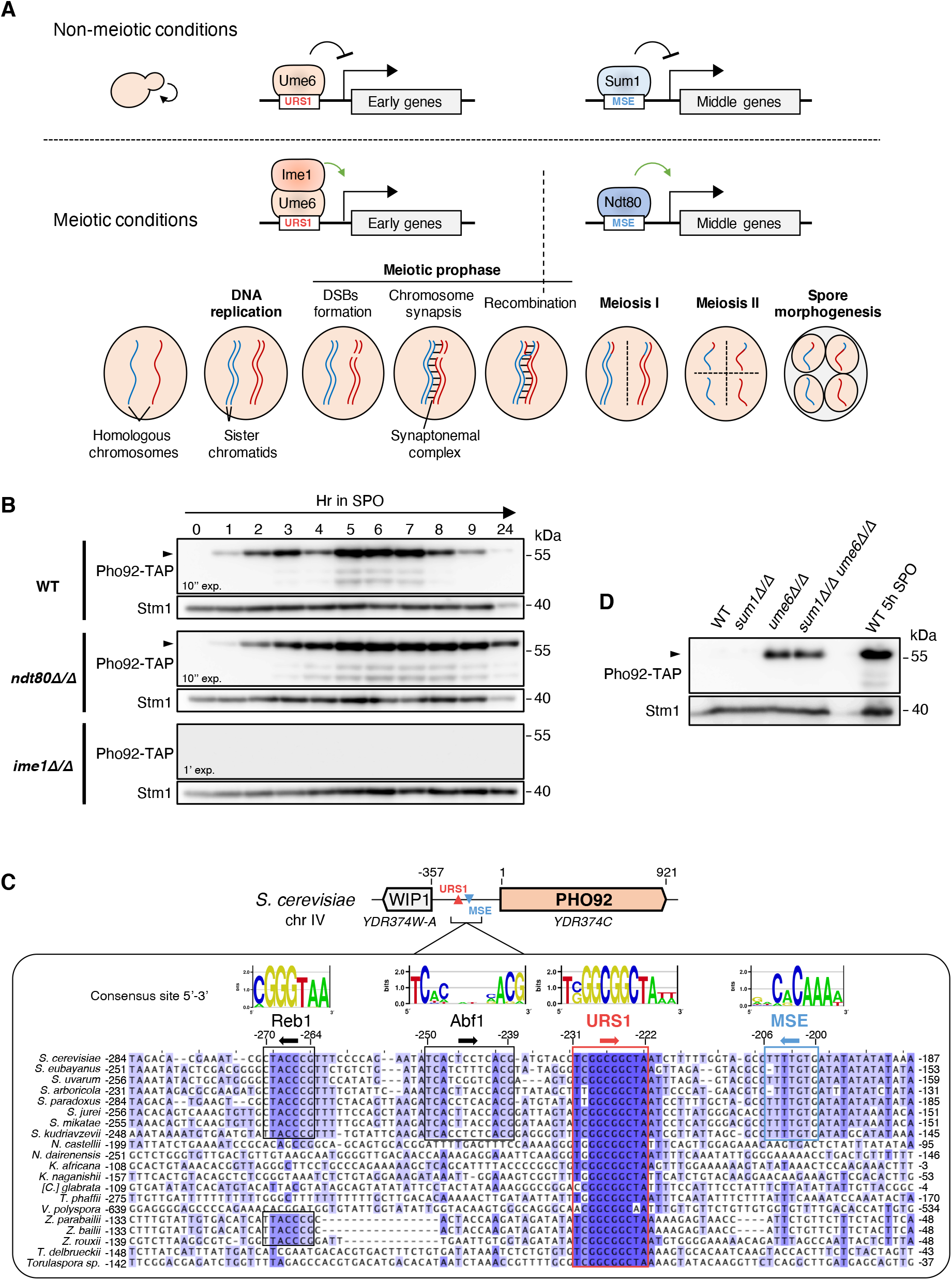
Pho92 is an early meiosis factor transcriptionally regulated in an Ime1/Ume6-dependent manner. (**A**) Schematic model of the transcriptional regulation of early and middle meiotic genes and the different stages of the meiotic program. Early genes are involved in activation of DNA replication and initiation of the meiotic prophase and are regulated by the presence of an URS1 element in their promoters, allowing their repression by Ume6 in non-meiotic conditions and their activation by Ime1 in meiotic conditions. Meiotic middle genes, involved in completion of the meiotic recombination, activation of meiotic divisions and spore formation, contained an MSE motif and are regulated in a Sum1- and Ndt80-dependent manner in non-meiotic and meiotic conditions, respectively. (**B**) Expression of Pho92-TAP during meiosis in diploid strains of the SK1 background. Western blots using Peroxidase Anti-Peroxidase complex to detect Pho92-TAP in total protein extracts from WT, *ndt80Δ/Δ* and *ime1Δ/Δ* cells collected at different hours following resuspension in SPO medium. Stm1 signal detected in the same extracts was used as loading control. (**C**) Sequence alignment of *PHO92* promoters in representative species of the *Saccharomycetaceae* family. The positions indicated are relative to the ATG of *PHO92*. The consensus binding site indicated for each motif or transcription factor is extracted from the YeTFaSCo database (de Boer and Hughes, 2012). (**D**) Expression of Pho92-TAP in diploid strains of the SK1 background grown in rich media. Western blots using Peroxidase Anti-Peroxidase complex to detect Pho92-TAP in total protein extracts from WT, *sum1Δ/Δ*, *ume6Δ/Δ* and *sum1Δ/Δ ume6Δ/Δ*cells collected in log phase cultures grown in rich media (YPDA). Stm1 signal detected in the same extracts was used as loading control.

Meiotic expression of the *IME4* gene encoding the m^6^A-methyltransferase is tightly regulated. Indeed, the production of an antisense haploid-specific transcript ensures that only diploid cells express Ime4 (Hongay et al., 2006; Gelfand et al., 2011; van Werven et al., 2012). Furthermore, *IME4* expression is under an Ime1 control and thus only induced in sporulation conditions (Chia and van Werven, 2016). While this regulatory scheme is general, the meiotic requirement for Ime4 in various yeast genetic background differs. A recent model ascribes this variability to the Rme1 (Regulator of meiosis) factor which acts as a transcriptional repressor of the master regulator *IME1* (Bushkin et al., 2019). Indeed, in strains with high levels of Rme1, it has been proposed that, during sporulation, Ime4 would methylate a specific site in the 3’UTR of *RME1* transcripts, leading to its destabilization and efficient *IME1* activation. Yet, if the contribution of m^6^A to meiosis is now well established, less progress has been made in understanding how these modifications are decoded and how individual sites could affect sporulation.

The molecular consequences of m^6^A have been mainly attributed to m^6^A readers: RNA binding proteins that specifically recognize the methylated A. The most-studied readers correspond to evolutionarily conserved YTH domain-containing proteins, represented by one member in yeast, up to 5 in mammals and more than 10 in plants. The 5 mammalian members are divided in 3 YTHDFs and 2 YTHDCs that impact in a specific or redundant manner several processes. YTHDC proteins were shown to contribute to the regulation of various processes, including nuclear functions such as pre-mRNA alternative splicing and nuclear export (Kasowitz et al., 2018; Roundtree et al., 2017; Xiao et al., 2016). In contrast, early studies indicated that YTHDFs readers have specialized functions in acceleration of mRNA decay (Du et al., 2016; Wang et al., 2014) and in enhancing translation efficiency (Hsu et al., 2017b; Mao et al., 2019; Wang et al., 2015). More recently, the 3 cytoplasmic human YTHDFs were shown to bind equivalently to m^6^A sites and to recruit the CCR4-NOT mRNA deadenylation complex to m^6^A-modified mRNAs (Du et al., 2016; Zaccara and Jaffrey, 2020). These and related results obtained on YTHDF orthologues in zebrafish and mouse (Kontur et al., 2020; Lasman et al., 2020) suggest a unified model in which YTHDFs readers have an evolutionarily conserved role in destabilizing their methylated targets while having limited direct impact on translation. The yeast *S. cerevisiae* encodes a single YTH reader: Pho92 (Phosphate metabolism 92), also known as Mrb1 (Methylated RNA-binding protein 1) (Kang et al., 2014; Schwartz et al., 2013), that belongs to the YTHDF group according to sequence similarity (Scutenaire et al., 2018). Structural analysis revealed that it binds to RNA surrounding m^6^A residues and recognizes the modified residue through an evolutionarily conserved pocket formed by the YTH domain (Theler et al., 2014; Xu et al., 2015). Reminiscent of other eukaryotic YTHDFs, an early study suggested that Pho92 interacts with the CCR4-NOT deadenylation complex to accelerate decay of mRNAs involved in phosphate metabolism (Kang et al., 2014). However, this function is unlikely imputable to m^6^A presence as these data have been obtained in haploid cells growing in rich media during which the *IME4* expression is repressed and no m^6^A is detectable (Gelfand et al., 2011; Hongay et al., 2006). Thus, although Pho92 is established as a putative m^6^A reader, its molecular impact on methylated transcripts remains unexplored.

Here we show that the m^6^A reader Pho92 is induced early in sporulation conditions and required for the proper timing of meiotic recombination and subsequent entry in meiotic divisions. By combining high-throughput RNA sequencing and mapping of Pho92 binding sites by UV-crosslinking and analysis of cDNA (CRAC), we show that Pho92 is recruited on specific mRNAs in a m^6^A-dependent manner and contributes to their downregulation during the meiotic prophase. Additionally, while functional m^6^A sites remained ill-defined, we provide here evidence that mutating individual m^6^A site in key Pho92 targets impairs the temporal control of cognate mRNAs expression and impacts meiosis. Thus, our analysis reveals the important role of specific mRNA methylation sites in controlling gene expression and cell fate.

## RESULTS

### PHO92: an early meiotic gene

Given the meiosis restricted accumulation of m^6^A in budding yeast, we first determined when *PHO92* is expressed. For this, we generated a C-terminal TAP-tagged version of Pho92 in the high sporulation efficiency SK1 background. Western blot analyses of protein samples collected during a meiotic time-course revealed that no signal is detected before transfer in sporulation medium (SPO) (Figure 1B). Pho92-TAP quickly accumulates after resuspension in SPO, reaching a maximum level after 5-6 hr, before decreasing at later time points. This profile, similar to those observed for *IME4* mRNA and m^6^A levels (Agarwala et al., 2012) and consistent with changes in the *PHO92* mRNA level (Primig et al., 2000; Schwartz et al., 2013), prompted us to look for regulatory elements controlling *PHO92* transcription. Strikingly, alignment of the *PHO92* promoter sequences from representative species of the *Saccharomycetaceae* family revealed the presence of a conserved element with similarity to URS1 (Figures 1C and S1A). A motif similar to MSE is also present but only in the *Saccharomyces* subfamily. This suggested potential *PHO92* expression regulation in meiotic conditions by Ime1 and Ndt80, respectively. To test this possibility, Pho92-TAP levels were monitored in *ime1*Δ/Δ and *ndt80*Δ/Δ strains following resuspension in SPO medium. While Pho92-TAP is similarly induced in the *ndt80Δ*/*Δ* and wild-type (WT), no signal was detected during the whole time-course in the *ime1Δ*/*Δ* mutant (Figure 1B). These data indicate that Ime1 is strictly required for *PHO92* induction, most likely through the highly conserved URS1 element in its promoter. In contrast, the MSE motif in the Pho92 promoter is unlikely functional as Ndt80 is not required for the initial induction of *PHO92*, consistent with *NDT80* being induced later, around 5 hr in SPO (Chu and Herskowitz, 1998). However, the Pho92-TAP signal persists longer in the *ndt80Δ/Δ* strain compared to WT (Figure 1B), likely as a consequence of the prophase arrest of the mutant, suggesting that Ndt80 activity (indirectly) participates in the down-regulation of Pho92-TAP at later steps of meiosis.

Given that many meiotic genes are repressed in rich media, we also assessed Pho92-TAP expression during vegetative growth. As expected, its expression was not detected in WT cells growing in rich media (Figure 1D). Similarly, Pho92-TAP is not detectable in absence of Sum1 that represses MSE-containing genes in non-meiotic conditions. However, deletion of Ume6, the key repressor of URS1-containing genes in non-meiotic conditions, allows Pho92-TAP expression in rich media. No additional expression level was detected in cells lacking both Sum1 and Ume6 indicating that Ume6 is the main repressor of *PHO92* expression in diploid and haploid cells growing in rich media (Figures 1D and S1B). Accordingly, recent data from ChIP-exo-seq (chromatin immunoprecipitation, exonuclease digestion and DNA sequencing) performed in haploid cells in rich media (Rossi et al., 2021) revealed a direct binding of Ume6 to the URS1 element of *PHO92* (Figure S1C). Altogether, these results indicate that the *PHO92* gene is regulated through a conserved URS1 element in its promoter, leading to Ume6-dependent repression in vegetative growth and early Ime1-dependent activation in meiotic conditions. Thus, the Pho92 protein is an early meiotic factor whose expression coincides with m^6^A presence.

### Pho92 is required for the proper kinetics of meiosis

Previous studies suggested that *pho92Δ*/*Δ* mutants have decreased sporulation efficiency or delayed meiosis (Deutschbauer et al., 2002; Schwartz et al., 2013). To pinpoint the role of Pho92 during meiosis, we quantitatively followed this process by microscopic observations of DAPI stained cells from *pho92Δ*/*Δ* and wild-type strains after transfer in SPO medium. To increase meiotic synchronicity, we used a derivative of the SK1 strain in which the promoter of the meiosis master regulator *IME1* is replaced by the inducible *CUP1* promoter (Chia and van Werven, 2016). This strain was used for all following experiments unless otherwise stated. In this background, where *IME1* expression is induced after 2 hr in SPO with addition of copper (II) sulfate, meiotic cells (characterized by > 1 nucleus) appear consistently ∼1 hr later in the *pho92Δ*/*Δ* strain compared to WT (Figure 2A). A longer delay ensues from the *ime4Δ*/*Δ* mutation used as control, consistent with previous observations (Schwartz et al., 2013). Monitoring the first and second meiotic divisions indicates that the delay results from defective entry in meiotic nuclear divisions. Indeed, in the absence of Pho92, bi-nucleated cells appear later but the kinetic progression between meiosis I and II, assessed by the appearance of tetra-nucleated cells, is not affected (Figure 2B). The overall sporulation efficiency of *pho92*Δ/Δ cells, monitored after 24 hr, was similar to WT but was reduced in the *ime4*Δ/Δ mutant (Figure S2A). Similar observations were made in the SK1 strain harboring the endogenous *IME1* promoter, even though with lower synchronicity (Figures S2B-C), demonstrating that the *pho92Δ*/*Δ* defect also arises upon endogenous *IME1* induction. We conclude that Pho92 is required for the proper kinetics of meiosis by regulating early event(s) prior to, or concomitant with, the first meiotic division and, at least in part, downstream of *IME1*. Interestingly, Pho92 inactivation does not phenocopy deletion of the Ime4 m^6^A methylase suggesting that m^6^A methylation also impacts processes independent of Pho92.

**Figure 2.**
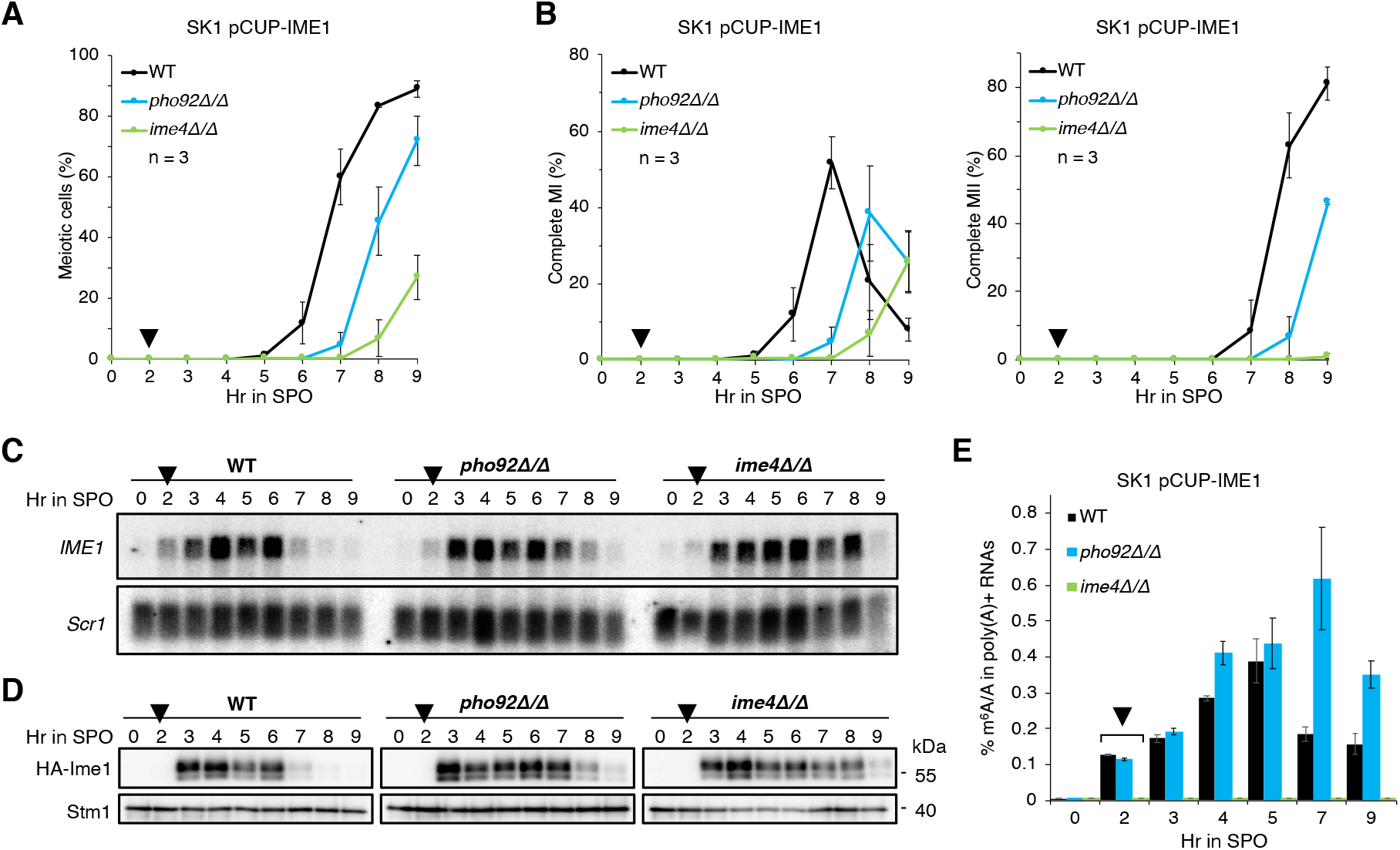
Pho92 is required for the proper timing of meiotic progression and impacts *IME1* and m^6^A down-regulation. (**A**) Kinetics of meiotic cells appearance in WT, *pho92Δ/Δ* and *ime4Δ/Δ* cells following resuspension in SPO medium and induction of *IME1* after 2 hr by addition of copper (II) sulfate. Meiotic cells correspond to cells containing > 1 nucleus as monitored by DAPI staining on 200 cells per time point. Means and standard deviations from 3 biological replicates. Black triangle above 2 hr: time of *IME1* induction. (**B**) Similar to (A) but with the kinetics of cells completing meiosis I (2 nuclei per cell; left panel) and meiosis II (> 2 nuclei per cell; right panel). Means and standard deviations from 3 biological replicates. Black triangle above 2 hr: time of *IME1* induction. (**C**) Northern blot analysis of *IME1* expression during meiosis in WT, *pho92Δ/Δ* and *ime4Δ/Δ* cells. The ncRNA ScR1 was used as loading control. Black triangle above 2 hr: time of *IME1* induction. (**D**) Western blot using anti-HA antibody to detect HA-Ime1 in total protein extracts from WT, *pho92Δ/Δ* and *ime4Δ/Δ* cells collected at different hours following resuspension in SPO medium. Stm1 signal detected in the same extracts was used as loading control. Black triangle above 2 hr: time of *IME1* induction. (**E**) LC-MS/MS quantification of m^6^A relative to A in polyadenylated RNAs extracted during a meiotic time course in two biological replicates of WT, *pho92Δ/Δ* and *ime4Δ/Δ* cells. No m^6^A was detected at 0-hr SPO nor at any time point of the *ime4Δ/Δ* mutant. Means, minimum and maximum measurements from 2 biological replicates. Black triangle above 2 hr: time of *IME1* induction.

### Pho92 impacts *IME1* and m^6^A down-regulations at late prophase

Having shown delayed meiosis upon deletion of *PHO92*, we tested whether the *IME1* expression was affected at RNA and protein levels in the pCUP-IME1 strains. Northern and western blot analyses using material extracted during sporulation time-courses revealed that *IME1* is strongly induced at mRNA and protein levels in the WT, *pho92*Δ/Δ and *ime4*Δ/Δ strains at 3 hr in SPO medium, corresponding to 1 hr after addition of copper (II) sulfate (Figures 2C-D). No *IME1* mRNA nor protein were detected without addition of copper at this time in WT cells (Figure S2D). Although the induction of *IME1* expression is similar between strains, its down-regulation occurs with different kinetics. In the WT, *IME1* mRNA and protein levels decrease after 6 hr in SPO medium, while they decrease after 7 hr in the *pho92Δ*/*Δ* mutant and after 8 hr in *ime4Δ*/*Δ* cells (Figures 2C-D). Hence, the timing of *IME1* down-regulation correlates with completion of meiosis I which differs for each strain (Figure 2A). It is important to note that in the pCUP-IME1 background, Rme1, the repressor of *IME1* suggested to be the main target of m^6^A regulation (Bushkin et al., 2019), has no impact on meiotic activation. Indeed, both the accumulation of the Ime1 protein and meiotic progression are similar in *rme1Δ*/*Δ* cells compared to WT (Figures S2E-F) indicating that m^6^A acts by other target sites. Overall, these results show that deletion of *PHO92* does not interfere with the initial accumulation of *IME1* at RNA and protein levels but rather hampers the pathway leading to its down-regulation at middle/late steps of meiosis.

We then monitored the impact of Pho92 on m^6^A accumulation during meiosis. The m^6^A/A ratio present in bulk poly(A) RNAs was quantified by Ultra-Performance Liquid Chromatography-tandem Mass spectrometer (UPLC-MS/MS). No m^6^A was detected in the entire time-course in the *ime4Δ*/*Δ* mutant, used as negative control (Figure 2E). In both WT and *pho92Δ*/*Δ* mutant, m^6^A starts to accumulate after 2 hr in SPO medium. Then, m^6^A levels reach a maximum after 5 hr in SPO medium in WT, corresponding to the onset of prophase I, and decrease afterwards. This is in agreement with data showing that mRNA methylation peaks during the end of G2/prophase and decreases following Ndt80 activation and the beginning of meiotic divisions (Agarwala et al., 2012; Schwartz et al., 2013). In the *pho92Δ*/*Δ* mutant, the accumulation of m^6^A continues, reaching a maximum at 7 hr, and only starts to diminish afterwards (Figure 2E). This suggests that methylated mRNAs are stabilized or more abundantly produced in the *pho92Δ*/*Δ* mutant at middle stages of meiosis. Combined with the prolonged expression of *IME1*, our data reveal that the *pho92Δ*/*Δ* mutation is likely extending the duration of the meiotic prophase by lengthening the persistence of methylated transcripts.

### Pho92 affects the meiotic transcriptome during prophase

To unravel the global impact of Pho92 on gene expression compared to Ime4 during meiosis, we collected samples at different time points (0, 2, 3, 5, 6, 7 and 9 hr) following SPO resuspension in biological triplicates of WT, *pho92Δ*/*Δ* and *ime4Δ*/*Δ* cells (Figures 3A & S3A). RNAs were extracted and poly(A) RNAs subjected to sequencing. We acquired a mean of > 22 million reads per sample and obtained normalized expression values (read count ⩾ 1 in at least one sample) on 6,092 genes, covering more than 92% of the latest version of the yeast transcriptome (6611 ORFs as of March 2022). Comparison of the sequencing data by principal component analysis (PCA) indicated that the 3 strains have highly related transcriptome profiles at the early time points (up to 3 hr) but start to strongly diverge from each other after 5 hr in SPO medium (Figure 3B). Yet, this analysis shows that differences are only transient as they follow similar trajectories with roughly a 1 hr delay for *pho92Δ*/*Δ* and a 2-3 hr delay for *ime4Δ*/*Δ*. Accordingly, the number of differentially expressed genes compared to WT (fold change ⩾ 2, adjusted p-value < 0.05) increases over time after 5 hr in SPO medium in both *pho92Δ*/*Δ* and *ime4Δ*/*Δ* cells (Figure S3B). In line with the more severe phenotype observed in *ime4Δ/Δ*, the number of differentially expressed genes at time points beyond 5 hr is between 1.4 and 2.4-fold more important in this strain compared to *pho92Δ*/*Δ*. Thus, the transcriptomic analysis reinforces our cellular and molecular analyses indicating that *pho92Δ*/*Δ* and *ime4Δ*/*Δ* defects become apparent around the first meiotic division, 5 hr after transfer to SPO medium.

**Figure 3.**
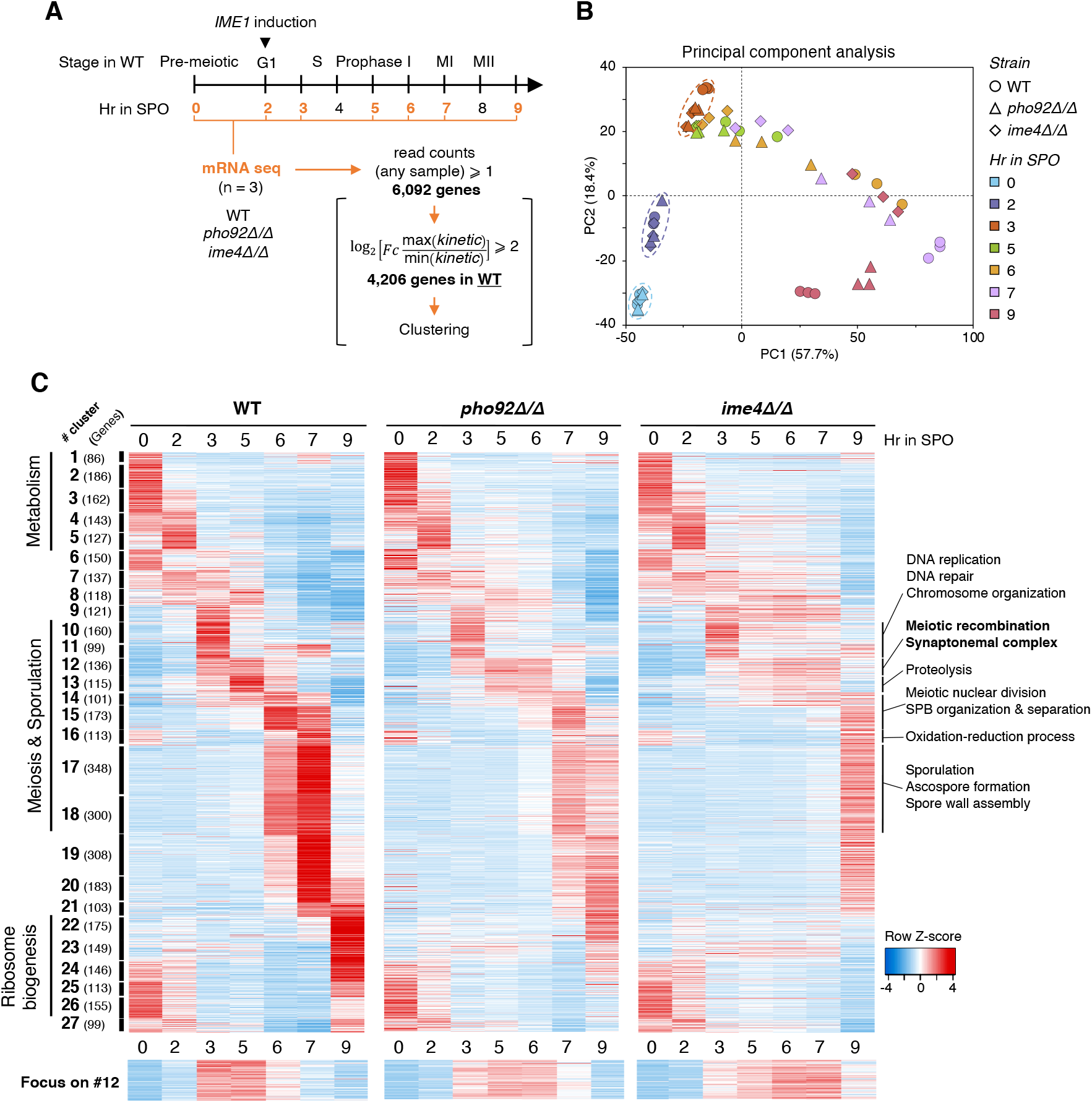
The meiotic transcriptome program is delayed in absence of Pho92. (**A**) Representation of collected samples for mRNA-seq during the meiotic course. The stage in WT cells is indicated above the time line. Time points in orange represent the ones used for mRNA-seq analyses. Below are the criteria used for the genes considered for further analyses and clustering data represented in (C). (**B**) Principal component analysis based on the normalized and stabilized gene expression levels of all samples in 3 biological replicates. The genotype of the strain is illustrated by the shape of the dot and colors represent the different time points across the meiotic time course. (**C**) Heat map representation of the 4,206 genes selected according to criteria in (A) following meiosis induction in WT cells, *pho92Δ/Δ* and *ime4Δ/Δ* mutants. Z-score transformation was performed for each gene considering the mean normalized read count between three biological replicates of each sample. Each row represents one gene and each column represents a sample. Z-score is scaled for each line from red (maximum expression value from all samples) to blue (lowest expression value from all samples). Genes with similar expression pattern were clustered and arranged together according to the chronology of their respective maximum expression in the WT strain. At the left is indicated the major biological processes enriched in different group of clusters, the number ID of each cluster (#) and, in parenthesis, the number of genes in each cluster. No known biological processes were significantly enriched in clusters #6 to #9 and #19 to #21. At the right, a more detailed list of biological processes is indicated for key meiotic clusters only. Lower panel represents a focus on cluster 12 enriched in genes involved in meiotic recombination and synaptonemal complexes structures.

To compare gene expression in the two mutants relatively to WT, we selected the 4,206 genes that present a fold change of at least 4 between their maximum and minimum expression values over the meiotic time-course in the WT (Figure 3A) and performed a clustering analysis creating 27 groups of expression profiles (Figure 3C). Plotting the expression data of these clusters in the *pho92Δ*/*Δ* and *ime4Δ*/*Δ* mutants reveals similar waves of expression from 0 to 5 hr. Indeed, transcripts of clusters #1-5 and #24-27, enriched in general metabolism and ribosome biogenesis-related biological processes respectively, are quickly down-regulated following transfer in SPO medium in all strains. Then, at 3 hr in SPO medium, mRNAs from genes in clusters #10-12 enriched in meiotic functions start accumulating. This includes genes with functions in DNA replication and repair, meiotic recombination and synaptonemal complexes structures. Interestingly, mRNAs corresponding to cluster #12 are strongly repressed at 6 hr in SPO in the WT but remain abundant at this stage in *pho92Δ*/*Δ* and *ime4Δ*/*Δ* cells (Figure 3C), spreading their peak of expression. At this time point, downstream clusters #14-18, enriched in factors involved in meiotic nuclear divisions and spore morphogenesis, are expressed in the WT strain but their induction is delayed and less prominent in the *pho92Δ*/*Δ* mutant and even more so in the *ime4Δ*/*Δ* strain. Thus, these data show that, Pho92 and Ime4 have a limited impact on the transcriptome early after transfer in SPO medium but rather affect processes at the onset of meiotic prophase. Interestingly, one of the first clusters affected by inactivation of *PHO92* and *IME4* contains transcripts encoding factors involved in meiotic recombination and synaptonemal complex structures (Figure 3C, lower panel).

### Pho92 temporally impacts the formation of meiotic DNA double-strand breaks

As our results indicate that the earliest detectable molecular defects resulting from *PHO92* and *IME4* deletions occur around prophase I, we first assessed whether Pho92 or Ime4 impact DNA replication. Flow-cytometry analysis showed that DNA replicates in similar manner between 3 and 4 hr after resuspension in SPO medium in the WT and the two mutants (Figure 4A). Consistently, our transcriptomic data indicate that the key genes involved in DNA replication are similarly induced at mRNA levels at 3 hr in SPO medium in all strains (Figure S4A). In contrast, most genes involved in meiotic recombination show an extended expression in *pho92Δ*/*Δ* and *ime4Δ*/*Δ* mutants compared to WT (Figure S4B), in accordance with their belonging to cluster #12 (Figure 3B). To assess the timing of meiotic recombination in the different strains, we monitored phosphorylation of Hop1, a known indirect marker of meiotic DSBs formation (Carballo et al., 2008; Niu et al., 2005), and of Hed1 at Tyr40, a target site of the Mek1 kinase active during the meiotic recombination checkpoint (Callender et al., 2016; Prugar et al., 2017). Western blot using specific antibodies show that both are mostly phosphorylated at 5 hr in SPO medium in the WT (Figure 4B). In contrast, in absence of Pho92 or Ime4, the phosphorylated forms peak at 6 hr and p-Hed1 remain detectable for few additional hours in the *ime4Δ*/*Δ* cells. These observations suggest a delay in meiotic DSBs formation as well as a slightly prolonged time of DSBs repair in the absence of Pho92, an effect that is even more pronounced in the absence of Ime4. To confirm this hypothesis, we directly determined the appearance of DSBs by Southern blot in a DSBs hot-spot near the *ARE1/YCR048w* gene (Baudat and Nicolas, 1997; Buhler et al., 2007) (Figures 4C-D). In this region, meiotic DSBs are visible at 4 and 5 hr in the WT and then disappear afterwards as a consequence of DSBs repair (Figure 4D). Consistent with data presented above, DSBs appear 1 hr later in *pho92Δ*/*Δ* and *ime4Δ*/*Δ* cells and remain visible for few additional hours. Interestingly, while the difference at late meiotic stages between *pho92Δ*/*Δ* and *ime4Δ*/*Δ* mutants is clear, the initial delay observed in DSBs formation is similar, suggesting that both factors have a comparable impact on meiotic progression until this stage. Thus, our results show that Pho92 and m^6^A presence are required for the proper timing of meiotic DSBs formation and in their absence, the meiotic recombination checkpoint, supported by Mek1 activity during DSBs repair, is likely more prolonged.

**Figure 4.**
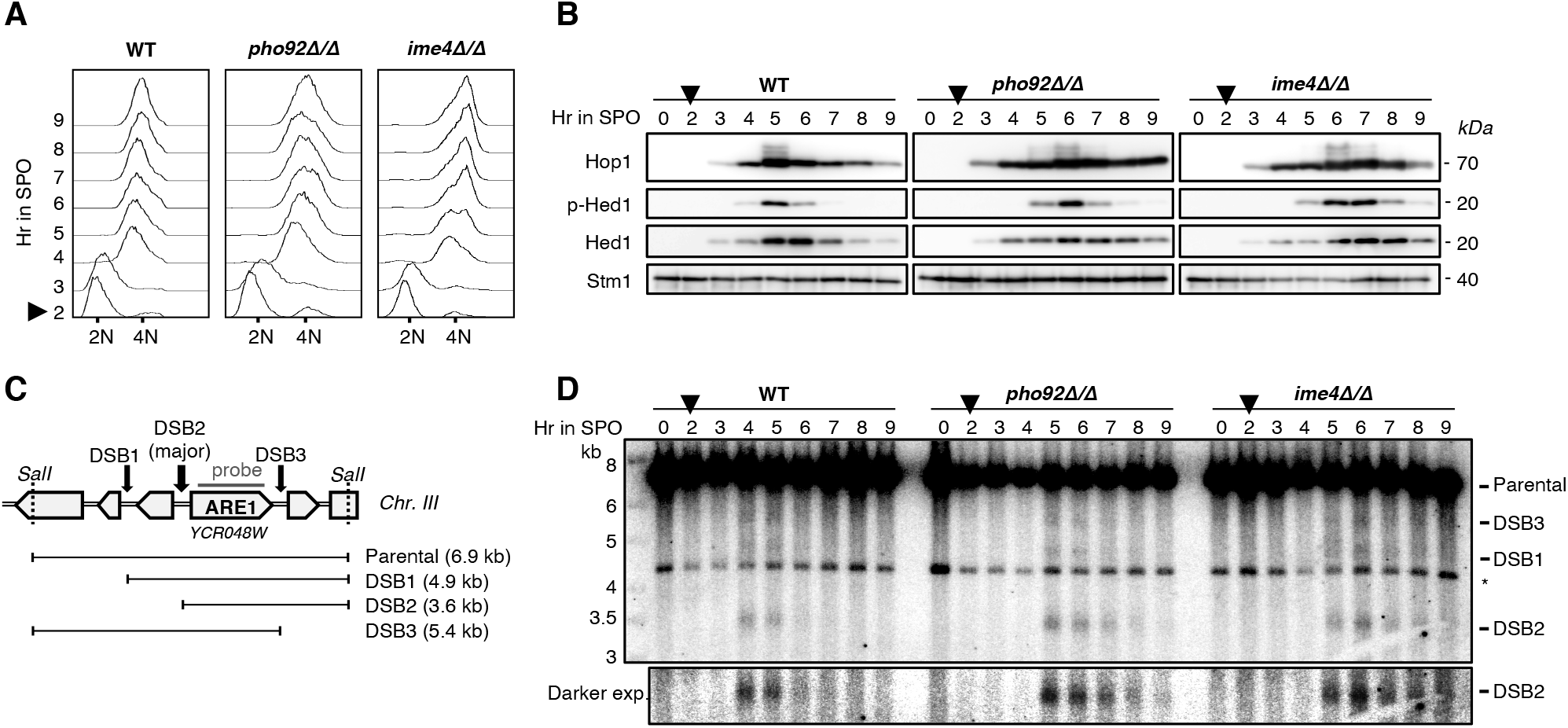
The absence of Pho92 delays meiosis after DNA replication but prior to DSB formation. (**A**) Flow cytometry analysis of DNA content in WT, *pho92Δ/Δ* and *ime4Δ/Δ* cells across a meiotic time course. Data were collected from cells harvested from 2 hr (time of *IME1* induction indicated by a black triangle) to 9 hr after resuspension in SPO. At least 10,000 single cells were counted for each time point. (**B**) Western blot analysis of Hop1, p-Hed1 (Hed1 phospho-Tyrosine 40) and Hed1 in total protein extracts from WT, *pho92Δ/Δ* and *ime4Δ/Δ* cells following resuspension in SPO medium. Stm1 signal detected in the same extracts was used as loading control. Black triangle above 2 hr: time of *IME1* induction. (**C**) Schematic representation of the *YCR048W* region from the chromosome III and the different fragments generated after meiotic DSBs formation and SalI enzymatic digestion. The position of the probe used to detect all fragments is indicated in dark grey solid line. (**D**) Southern blot analysis of meiotic DSBs formation in the *YCR048W* region in WT, *pho92Δ/Δ*and *ime4Δ/Δ* cells following resuspension SPO medium. An unspecific fragment (*) of ∼4.3 kb is detected from cross reaction of the radiolabeled probe with a SalI-generated fragment of the ARE2/*YNR019W* gene, paralogous to ARE1/*YCR048W*. Lower panel represents a focus with darker exposure on the fragment generated by the DSB2, as illustrated in (C). Black triangle above 2 hr: time of *IME1* induction.

### Pho92 directly binds methylated mRNAs involved in meiotic recombination

To identify m^6^A-dependent mRNA targets bound by Pho92 during meiotic recombination, we performed an improved version of the UV-Crosslinking and analysis of cDNAs (CRAC) technique (Candelli et al., 2018; Granneman et al., 2009; Villa et al., 2020) on samples harvested at 5 hr in SPO medium. As required for the procedure, we used cells expressing an HTP (His_6_-TEV-ProteinA)-tagged version of Pho92 in WT and in *ime4Δ*/*Δ* strain as a control, to assess its specific binding to m^6^A-modified RNAs. Pho92-HTP cells without UV-crosslinking and a wild-type extract, in which no HTP-tagged protein is expressed, were also used as controls. We first confirmed the functionality of the Pho92-HTP protein by verifying that meiosis progression is not delayed in cells harboring this fusion and that the tagged protein accumulates with the same profile as the TAP-tagged version following *IME1* induction, in both the WT and *ime4Δ*/*Δ* backgrounds (Figures S5A-B). Following the CRAC procedure, binding sites were identified in reads presenting a specific deletion induced at UV-crosslinked sites by the reverse-transcriptase reaction during the library preparation. To compare CRAC data between different samples and replicates, an internal spike-in was used consisting of UV-crosslink cells of *Schizosaccharomyces pombe* expressing the essential RNA-binding protein Luc7 fused to the HTP tag (Figure S5C). The binding score to the Luc7 target, U1 RNA, was used to normalize the scores of peaks identified in *S. cerevisiae* (Figure S5D). After normalization, the highest numbers of peaks (score > 0.9) were found in the two biological replicates of Pho92-HTP, as expected, with 1061 and 1101 peaks respectively (Figure S5E). Considering the 794 peaks commonly found in the replicates of Pho92-HTP, we classified them according to their scores in the other conditions. We found 627 Ime4-dependent peaks that we further divided in two groups: the 295 peaks with high scores (> 4; corresponding to scores above the 99.2 percentile of the distribution of all peaks) in both replicates of Pho92-HTP and at least 2-fold higher than in the *ime4Δ*/*Δ* mutant (Figure S5F), were selected as “high-score” Ime4-dependent binding sites of Pho92 while the 332 remaining peaks (score < 4 in one replicate of Pho2-HTP) were considered as “lower-score” (Figures 5A). In other control samples, no peaks or peaks with very low score were observed at these sites. The remaining peaks were defined as “Ime4-independent” binding sites (142 peaks), when they present scores > 2 in *ime4Δ/Δ* mutant, and “unspecific” sites (25 peaks) with score > 2 in Pho92-HTP no UV and/or WT cells (Figure 5A). The “high-” and “lower-score” Pho92-binding sites show similar profiles with 259 peaks (87.7%) and 293 peaks (88.2%) localizing in CDS and predicted UTR regions of 208 and 253 mRNAs, respectively (Figure 5B). The majority of the mRNA targets harbor one Pho92-binding site but some have up to 7 sites (Figure S5G). In contrast, most, if not all, peaks of the “Ime4-independent” and “unspecific” groups are found in tRNAs and rRNAs, thus likely reflecting false positive signals. Strikingly, a single motif was significantly enriched in Pho92-binding sites corresponding to the consensus AWRGACDWU (W: A or U, R: A or G, D: A, G or U). It was found in 286 (96,9%) and 324 (97,6%) peaks of the “high-score” and “lower-score” binding sites respectively, and it matches to an extended version of the established m^6^A methylation consensus site (Figures 5C and S5H). Notably, the position +4 following the central A is strongly biased toward an “U” residue, found in 97% and 93% of the cases in “high-score” and “lower-score” binding sites, respectively. Sequence analysis under the peaks of the other populations did not reveal similar motifs (Figure S5H). In addition, considering only the peaks identified in mRNAs, binding of Pho92 shows a bias toward the 3’ end of coding sequences surrounding the stop codon (Figure 5D). These findings are fully consistent with previous analyses of m^6^A sites signature arguing that Pho92 binds Ime4-methylated mRNAs. However, 98 targets (47%) and 121 targets (47.8%) of the “high-score” and “lower-score” populations, respectively, were not identified as methylated candidates in the previous m^6^A-seq (Schwartz et al., 2013) or MAZTER-Seq (Garcia-Campos et al., 2019; group confidence > 1) analyses (Figure S5I), possibly owing to strain or condition differences. Thus, our approach, that directly target Ime4-dependent binding sites of Pho92, reveals the existence of new populations of methylated mRNAs during yeast meiosis.

**Figure 5.**
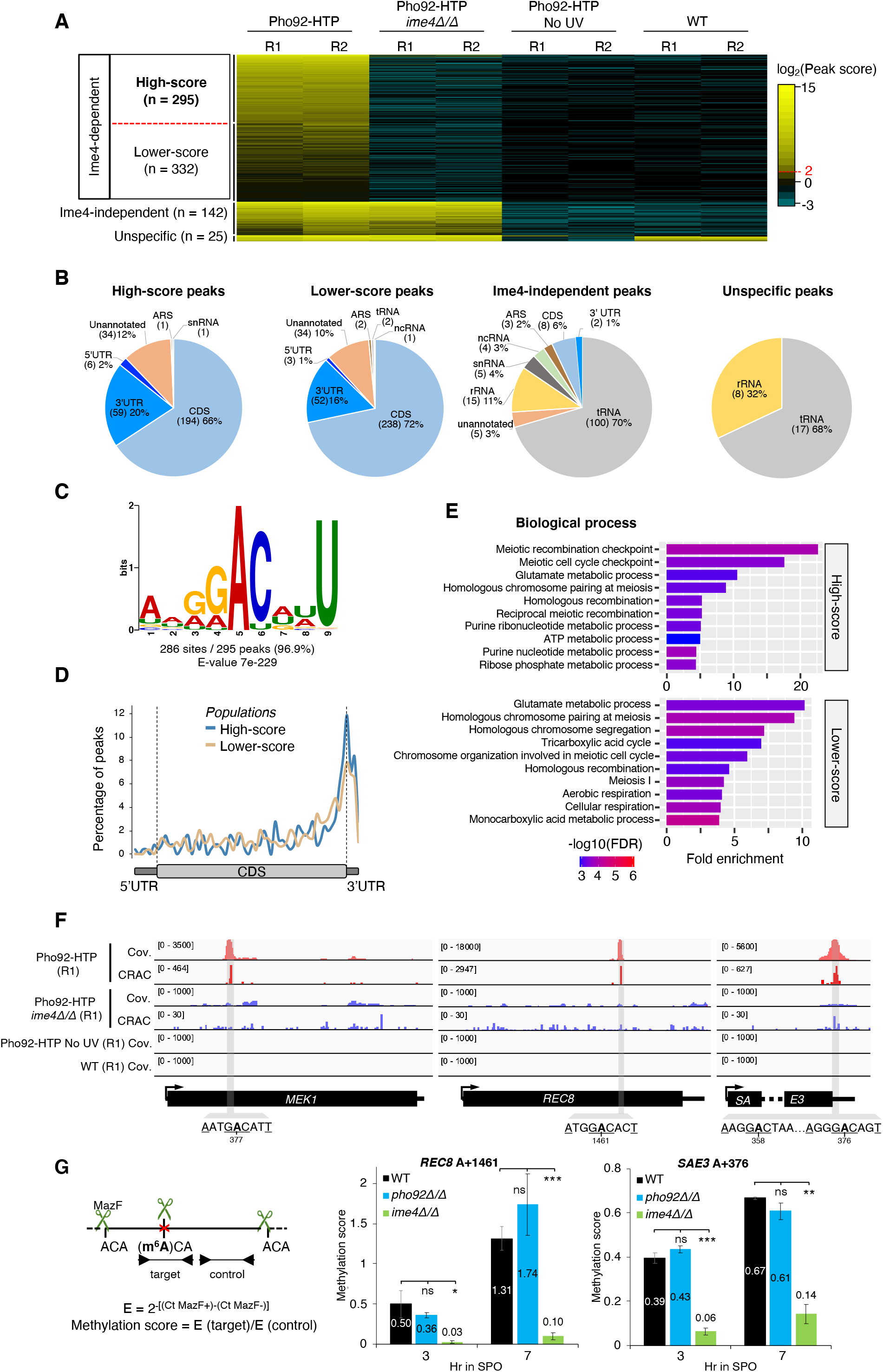
The mRNA targets of Pho92 identified by CRAC are mostly bound in an Ime4-dependent manner. (**A**) Heat map showing the log2-transformed peak scores for the 794 peaks commonly found in the two biological replicates of Pho92-HTP (scores > 0.9) in the CRAC datasets and their respective values in other conditions (*ime4Δ/Δ* mutant, No UV-crosslinking, WT with no HTP expression). Peaks were clustered and arranged together according to the fold change score (Pho92-HTP/Pho92-HTP in *ime4Δ/Δ*). (**B**) Pie charts showing peak distribution in different types of RNA and mRNA regions in the different defined peak populations. (**C**) Sequence motif of the Pho92-binding sites as determined by MEME, based on the sequences under the 295 high-score Ime4-dependent peaks. (**D**) Metagene analysis of the Pho92-binding sites based on the peaks’ summit positions of the high-score Ime4-dependent population (blue line) and lower-score Ime4-dependent populations (light brown line). (**E**) Gene ontology enrichment analysis for Pho92-bound transcripts of the high-score (upper panel) and lower-score (lower panel) Ime4-dependent peak populations. Only the top 10 biological processes are shown and ranked according to the fold-enrichment against the genomic background. The -log10 (False Discovery Rate) value of each category is calculated based on nominal p-value from the hypergeometric test and indicated in a color scale from max (red) to lowest (blue) values. (**F**) IGV tracks showing specific Ime4-dependent binding of Pho92-HTP on *MEK1* (left), *REC8* (middle) and *SAE3* (right) transcripts. Shown are the sequence coverage (Cov.) for each sample and the crosslink signal (CRAC) detected by the nucleotide deletion induced at the UV-crosslinked binding sites. Only values from one biological replicate are shown for each condition (Pho92-HTP, Pho92-HTP *ime4Δ/Δ*, Pho92-HTP No UV, WT). The identified Pho92-binding peaks are highlighted in grey with the consensus binding sequences indicated at the bottom and the position of the central “A” relative to the ATG of each gene. (**G**) Determination of methylation scores by MazF-qPCR of two positions in the identified Ime4-dependent binding sites of Pho92 in *REC8* (middle) and *SAE3* (right) transcripts. Samples from WT, *pho92Δ/Δ* and *ime4Δ/Δ* cells were taken at 3 and 7 hr following resuspension in SPO medium in which *IME1* was induced after 2 hr. Means and standard deviations from 3 biological replicates. ns: p > 0.05, *p < 0.05, **p < 0.01, ***p < 0.001, two-tailed *t-*test compared to WT samples at the considered time point. Left panel: schematic representation of the MazF-qPCR assay. The methylation scores are estimated by the abundance of the “target” amplicon in MazF-treated versus untreated samples, normalized to the abundance of the “control” amplicon in MazF-treated versus untreated samples.

Gene ontology (GO) analysis revealed that the Pho92 targets are highly enriched in meiotic recombination functions, chromosome pairing, meiosis I and also diverse metabolic processes (Figure 5E). Interestingly, the former group correlates with the first cluster found to be affected in *pho92Δ/Δ* cells in our transcriptomic data (Figure 3C). In the high-score population, the set of 16 genes with meiotic functions comprises major actors involved in meiotic checkpoint (*IME2*, *MEK1, PCH2, DDC1, RAD6*), members of the cohesin and synaptonemal complexes (*REC8, SMC3, RED1, HFM1, HOP1, HOP2)* and components of the meiotic recombinase (*DMC1, SAE3, MEI5*) (examples in Figures 5F and S5J). To independently confirm the methylation status of some target sites in a more quantitative manner, we set up MazF-qPCR assays, a recently developed method that allows interrogating methylation levels at a subset of sites with m^6^ACA sequence (Garcia-Campos et al., 2019). This confirmed Ime4-dependent methylations at the Pho92-binding sites on the transcripts of *REC8* at position A+1461 (relative to A of ATG in genomic DNA) and *SAE3* at position A+376, at 3 and 7 hr after transfer in SPO medium (Figure 5G). Notably, these sites were not found in m^6^A seq (Schwartz et al., 2013) but were identified in low confidence sites (group 1) in the MAZTER-seq data (Garcia-Campos et al., 2019). These data indicate that Pho92 is directly binding mRNAs during meiotic prophase and that, most if not all of its targets, in particular those encoding meiotic recombination factors, are m^6^A-modified at the binding sites.

### Pho92 contributes to the down-regulation of its meiotic targets

To establish the molecular consequences of RNA binding by Pho92, we correlated profiles of mRNA levels determined by RNA-seq data during meiosis with Ime4-dependent targets of Pho92. Even if the previously defined “high-score” and “lower-score” populations showed similar features, we only considered hereafter the set of “high-score” targets to strengthen our conclusions. We first observed that the 208 Pho92 mRNA targets have a higher average expression compared to all genes expressed at 5 hr in SPO, the time in which CRAC data were obtained (Figure 6A). Indeed, 148 targets (71%) are among the top 25% of genes with higher expression values at 5 hr in SPO in all strains, suggesting that Pho92 affects a significant proportion of the transcriptome.

**Figure 6.**
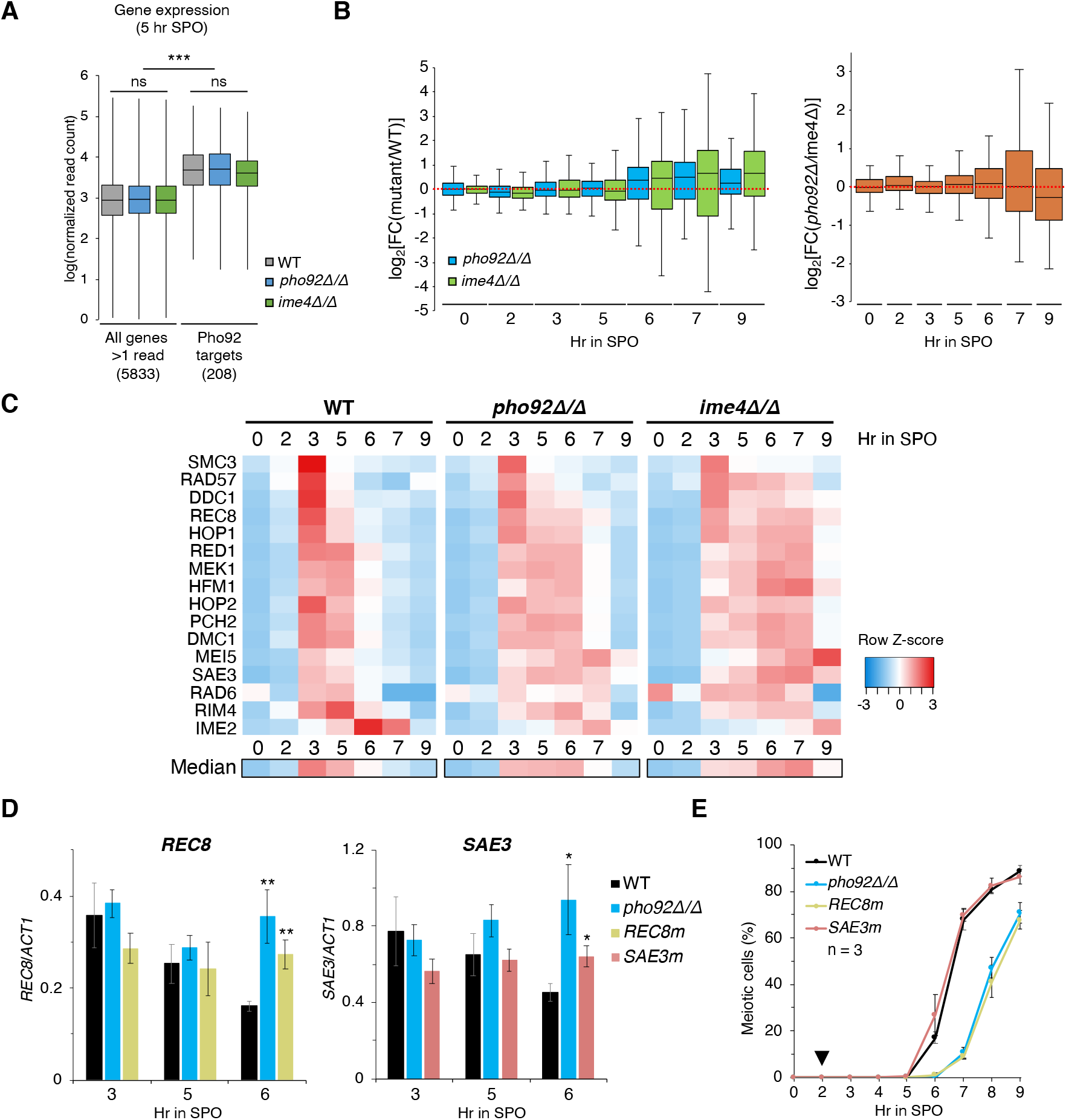
Pho92 contributes to the down-regulation of its meiotic targets including *REC8* and *SAE3*. (**A**) Expression of the Pho92-targets compared to all genes expressed (read >1) at 5 hr in SPO in WT, *pho92Δ/Δ* and *ime4Δ/Δ* cells. The distribution of the log10 normalized read counts of all genes is indicated with box plots (min, first quartile, median, third quartile, max) according to the mean values from 3 biological replicates. ns: p > 0.05, ***p < 0.001, two-tailed *t-*test. (**B**) Differential analysis of expression of Pho92 targets in *pho92Δ/Δ* and *ime4Δ/Δ* mutants compared to WT in mRNA-seq data. Left panel: shown are the log2 fold change value of the mean normalized read count of 3 biological replicates of each Pho92 target in *pho92Δ/Δ* mutant (blue) or *ime4Δ/Δ* mutant (green) over the WT across the meiotic time course. Right panel: expression values of the Pho92 targets in the *pho92Δ/Δ* mutant compared to the *ime4Δ/Δ* mutant. (**C**) Heat map representation of expression of the 16 Pho92 targets with established meiotic functions during the meiotic time course in WT cells, *pho92Δ/Δ* and *ime4Δ/Δ* mutants. Z-score transformation was performed for each gene considering the mean normalized read count of three biological replicates. Each row represents one gene and each column represents a sample. Z-score is scaled for each line from red (maximum expression value from all samples) to blue (lowest expression value from all samples). Genes are arranged together according to the chronology of their respective maximum expression in the WT strain. A median expression profile from the 16 genes is represented below. (**D**) RT-qPCR analysis of *REC8* (left) and *SAE3* (right) transcripts at 3, 5 and 6 hr after SPO resuspension in WT, *pho92Δ/Δ* and respective point mutation strains. *REC8m* indicates a point mutation of A+1461>G in *REC8 while SAE3m* indicates the point mutations C+359>T and A+376>C in *SAE3*. Means and standard deviations from 3 biological replicates. *p < 0.05, **p < 0.01, two-tailed *t-*test compared to the WT at each time point. (**E**) Kinetics of meiotic divisions from WT, *pho92Δ/Δ* mutant, *REC8m* and *SAE3m* point mutants used in (D). Meiotic cells correspond to cells containing >1 nucleus as monitored by DAPI staining on 200 cells per time point. Means and standard deviations from 3 biological replicates. Black triangle above 2 hr: time of *IME1* induction.

Our analysis of m^6^A levels and transcriptomic data during meiosis suggest a role for Pho92 in down-regulation of methylated mRNAs at a certain stage of meiosis. Accordingly, the Pho92 mRNAs targets accumulate to higher levels at 6, 7 and 9hr in SPO in *pho92Δ/Δ*compared to the WT (Figures 6B and S6A). At 6hr in SPO for example, 47 of the 208 Pho92 targets are significantly up-regulated in *pho92Δ/Δ* compared to the WT (Figure S6B). A similar effect is seen in the absence of *IME4*, even though with a larger variation in expression values and an increased differential expression compared to the WT (Figures 6B and S6A-B). Yet, closer inspection of mRNA profiles of the Pho92 targets during the meiotic time course revealed some temporal heterogeneity. In line with a delayed down-regulation, some targets, reaching their maximum mRNA level at 5 hr in SPO in the WT, accumulates to a higher level at 6 hr in absence of Pho92 (Figure S6C). However, other targets with peaks of expression at 6 and 7 hr in SPO in the WT, show a delayed up-regulation in *pho92Δ/Δ* cells (Figure S6C). The delayed up-regulation of this second group in absence of Pho92 (or Ime4) is likely an indirect consequence of the dysregulation of earlier phases of meiosis, although we cannot exclude that Pho92 has opposite effects on the temporal expression of different groups of targets. Interestingly, almost all of the 16 targets with functions related to meiotic recombination show similar profiles with an extended expression in *pho92Δ/Δ* and *ime4Δ/Δ* compared to WT (Figure 6C). We conclude that Pho92 is specifically required for the down-regulation of its mRNA targets reaching their maximum expression during meiotic prophase that include, in particular, genes associated with meiotic recombination.

### Altering m^6^A sites prevents normal down-regulation of Pho92 targets and impacts meiosis

To investigate whether m^6^A controls the fate of Pho92 meiotic mRNA targets, we introduced point mutations in the two Pho92-binding sites which we confirmed to be methylated by 2 independent strategies, namely sites in the Rec8 and Sae3 mRNAs (Figures 5F-G). The mutation A+1461>G was introduced in *REC8* while both C+359>T and A+376>C substitutions were introduced in two close sites in *SAE3*. In both of these cases, point mutation(s) altered the consensus methylation sequences without changing the encoded protein. The impact of these mutations on accumulation of the corresponding mRNA during meiosis was then assessed by RT-qPCR. In both cases, absence of Pho92 or mutation of the Pho92-binding sites does not alter the amount of *REC8* and *SAE3* mRNAs at 3 and 5 hr in SPO (Figure 6D). However, at 6 hr, the presence of point mutations abolishes the reduction of *REC8* and *SAE3* mRNA levels observed in WT cells, similarly to the defect observed in the *pho92Δ/Δ* mutant (Figure 6D). This observation prompted us to assay the kinetics of meiosis in these mutant strains. While mutation of the Pho92-binding sites in *SAE3* did not induce a change in the temporal appearance of meiotic cells compared to WT, the point mutant of *REC8* induced a meiotic delayed phenotype similar to the one observed in *pho92Δ/Δ* cells (Figure 6E). Hence, impairing the binding of Pho92 to *REC8* and *SAE3* mRNAs is sufficient to delay their down-regulation, and in the case of *REC8*, it also suffices to delay meiotic divisions, similar to the effects observed in the absence of Pho92. These results establish *REC8* as a critical target of m^6^A and Pho92 during meiosis.

## DISCUSSION

Modification of mRNAs through m^6^A-methylation has an evolutionarily conserved impact in gametogenesis of mammals (Hsu et al., 2017b; Wojtas et al., 2017; Xu et al., 2017; Zheng et al., 2013), flies (Hongay and Orr-Weaver, 2011), plants (Zhang et al., 2019) and yeast (Agarwala et al., 2012; Schwartz et al., 2013). However, the contribution of m^6^A reader as well as the relevance of individual m^6^A sites in mRNAs remain poorly known, in particular during yeast meiosis. Here we characterized the yeast m^6^A reader, providing an extended transcriptomic analysis across a synchronous meiosis and identifying direct, Ime4-dependent, targets of Pho92. We provide evidence that Pho92 controls the duration of the meiotic recombination and contributes to the regulation of transcript abundance, particularly of mRNAs encoding meiotic recombination factors. Through the direct analysis of sites bound by Pho92, we also shed light on functional sites contributing to the meiotic progression in yeast, among which *REC8* appears as a critical target.

A previous study suggested that Pho92 could regulate the stability of some transcripts in a phosphate dependent-manner in haploid cells (Kang et al., 2014). Our analysis casts some doubts on this function as Pho92 is not expressed in vegetative conditions and m^6^A is absent. *PHO92* expression analyses indicate rather that it is an early meiosis factor under the control of a URS1 element in its promoter, allowing both meiotic activation and repression in non-meiotic conditions. Interestingly, the *PHO92* promoter also contains a conserved binding-site for the general transcription activator Abf1 in *Saccharomyces* yeast. The association of an Abf1-binding site and an URS1 element is also found in promoters of *HOP1, RED1* and *ZIP1* (Gailus-Durner et al., 1996), 3 actors with specific functions in meiotic recombination. Altogether, these data support a specialized function of Pho92 during early meiotic steps.

Consistently, we observe that deletion of *PHO92* results in a delay in meiosis and of *IME1* downregulation with a concurrent dysregulation of the transcriptome profile during meiosis. Surprisingly, the absence of *PHO92* does not phenocopy the deletion of *IME4*, both for meiosis progression and for their impact on the transcriptome, the latter resulting in a more severe lag. Several models can account for these differences. First, Ime4 may have a function independent of its methylase activity, a possibility supported by the less severe phenotype induced by the catalytic site mutant of Ime4 (Agarwala et al., 2012). Alternatively, other reader protein(s) may be present during yeast meiosis or m^6^A may act by preventing the binding of some RNA binding proteins or by altering the structure of RNA. These possibilities are not mutually exclusive and further analyses will be required to pinpoint the mechanisms implicated. If the impact of *PHO92* and *IME4* deletions on meiosis differ, our kinetic analysis of transcriptome dynamics reveals very little differences during the first 3 hours of this process (Figures 3B and C). Starting at 5 hr in SPO, a lag is apparent in the *pho92Δ/Δ* mutant, a phenotype further exacerbated in the absence of Ime4. Yet, except for these delays, the same waves of transcripts are up- and down-regulated in WT and the two mutants, suggesting that a key checkpoint must be overcome to allow cells to progress further in meiosis and arguing against a global dysregulation of gene expression. Notably, the first group of genes with extended expression in the two mutants encode factors involved in meiotic recombination. This observation is consistent with the early steps of meiosis, including DNA replication, proceeding similarly in the 3 strains but DSBs formation, corresponding to the initiation of DNA recombination, occurring later in the two mutant strains, with little difference between them. Overall, our data indicate that the early stages of meiosis before the onset of meiotic prophase I are unaffected in the absence of m^6^A or of the m^6^A reader Pho92, but the absence of one or the other impacts meiotic recombination and delays downstream events. Strikingly, meiotic defects start to arise when or early after m^6^A peaks in WT cells.

Large sets of m^6^A sites had been previously reported in yeast mRNAs during meiosis (Bushkin et al., 2019; Dierks et al., 2021; Garcia-Campos et al., 2019; Leger et al., 2021; Schwartz et al., 2013). Those were established using different methods and show limited overlap, partly owing to distinct experimental parameters (different strains or analyses at different time points) but also possibly due to methodological limitations (McIntyre et al., 2020). Despite this large number of m^6^A sites, the evidence of functional sites able to affect gene expression at the molecular level and/or contributing to physiological function remain elusive. Until now, only the m^6^A site of the *RME1* transcript was reported to have an effect leading to its destabilization and early *IME1* activation (Bushkin et al., 2019). It is important to note, however, that *RME1* is repressed in diploid cells (van Werven and Amon, 2011) and that the impact of Ime4 on its expression was observed several hours after entry in sporulation and initial Ime1 expression (Bushkin et al., 2019), suggesting that the proposed regulation would at best affect late stage of the yeast meiotic program. Our observations question this model as our data argue that Pho92 functions downstream of *IME1*: first, we find no detectable levels of Pho92 protein in meiotic conditions in *ime1Δ/Δ* mutant cells; second, m^6^A levels increase concomitantly with Ime1 accumulation as a result of *IME4* induction; and third, the meiotic delay resulting from Pho92 inactivation is detected even after *IME1* induction in the pCUP-IME1 background, in which as anticipated, the absence of *RME1* has no impact. If our data do not support a role for Pho92 in *IME1* activation, we noticed that Pho92 influences the timing of *IME1* down-regulation, most likely indirectly as no Pho92 binding to the *IME1* mRNA was identified. Whether this a cause or a consequence for the delayed meiotic divisions in *pho92Δ/Δ* cells remains to be clarified.

To identify functional sites bound by Pho92 in an Ime4-dependent manner during yeast meiosis, we performed a CRAC analysis, a strategy that bypass possible biases associated with immuno-enrichment of m^6^A residues (Granneman et al., 2009). The CRAC data appeared relatively robust, with a good signal to noise ratio and with most Pho92 mRNA targets being clearly Ime4-dependent. At least in the cases of *SAE3* and *REC8* transcripts, the presence of methylated residues at different time points could be confirmed by an independent MazF-qPCR assay. If one could have envisaged that Pho92 binds non-methylated mRNAs, our CRAC data argue against this. Indeed, in the absence of Ime4, Pho92 fails to bind to its target mRNAs even though those are expressed, and “Ime4-independent” Pho92 binding sites appear to be nearly exclusively in non-coding RNAs. Yet, our Pho92 CRAC analysis identified significantly less modified mRNAs compared to previous analyses. Only 18-45% of Pho92-binding sites revealed by our study were also reported as methylated in 2 methodologically independent studies (Figure S5J). While some biological parameters (e.g., time point during meiosis or strain) may contribute to these differences, they could also be explained if the Pho92 reader only binds a subset of the methylated A residues, a possibility that will require further investigation.

It is noteworthy that the absence of Pho92 results in a prolonged presence of m^6^A and delayed down-regulation of the first groups of transcripts whose profiles are altered. Consistent with previous studies linking m^6^A readers with mRNA decay (Du et al., 2016; Zaccara and Jaffrey, 2020), we propose that Pho92 facilitates prophase exit by favoring down-regulation of key transcripts, notably those involved in meiotic recombination (Figure 7). Yet, the functional link between Pho92 and the decay machinery remains to be clarified. The identified Ime4-dependent targets of Pho92 include *REC8*, *HOP1* and *RED1*, encoding the core components of axial elements of the synaptonemal complexes that link homologous pairs of sister chromatids during meiotic recombination (Zickler and Kleckner, 2015). Rec8 is a meiosis specific member of the cohesin complex with major functions in cohesion of sister chromatids, formation of chromosome axis and promotion of reciprocal recombination during meiosis (Brar et al., 2009; Klein et al., 1999; Yoon et al., 2016). Alteration of either expression or clearance of Rec8 protein levels during meiosis impairs the beginning of the meiotic divisions (Brar et al., 2006, 2009). Other meiotic mRNA targets of Pho92 comprise *MEK1* and *SAE3*. Mek1 activity is linked to DSBs formation and is required for the meiotic recombination checkpoint (Prugar et al., 2017; Wu et al., 2010) while Sae3 is needed for full assembly of the synaptonemal complexes and DSBs reparation (Hayase et al., 2004; Mckee and Kleckner, 1997). Given that these studies showed an impaired meiotic prophase arrest when the expression of *SAE3* or *MEK1* are altered, this further support that Pho92 controls transcript abundance of key actors in the establishment of meiotic checkpoints. Surprisingly, we also observed, by MazF-qPCR assays, that *REC8* and *SAE3* mRNAs are methylated at 3 hr in SPO when no difference in the steady state levels of theses transcripts was observed between WT and *pho92Δ/Δ* mutant. This suggest that transcript methylation, at least on these two targets, is not temporally controlled and could indicate that the m^6^A impact on transcript degradation through Pho92 is regulated. One hypothesis that we did not explore is whether Pho92 could promote translation of its targets before facilitating their downregulation, in agreement with the presence of m^6^A in translating ribosomes (Bodi et al., 2015) and described functions of YTHDF readers in mammals (Wang et al., 2015).

**Figure 7.**
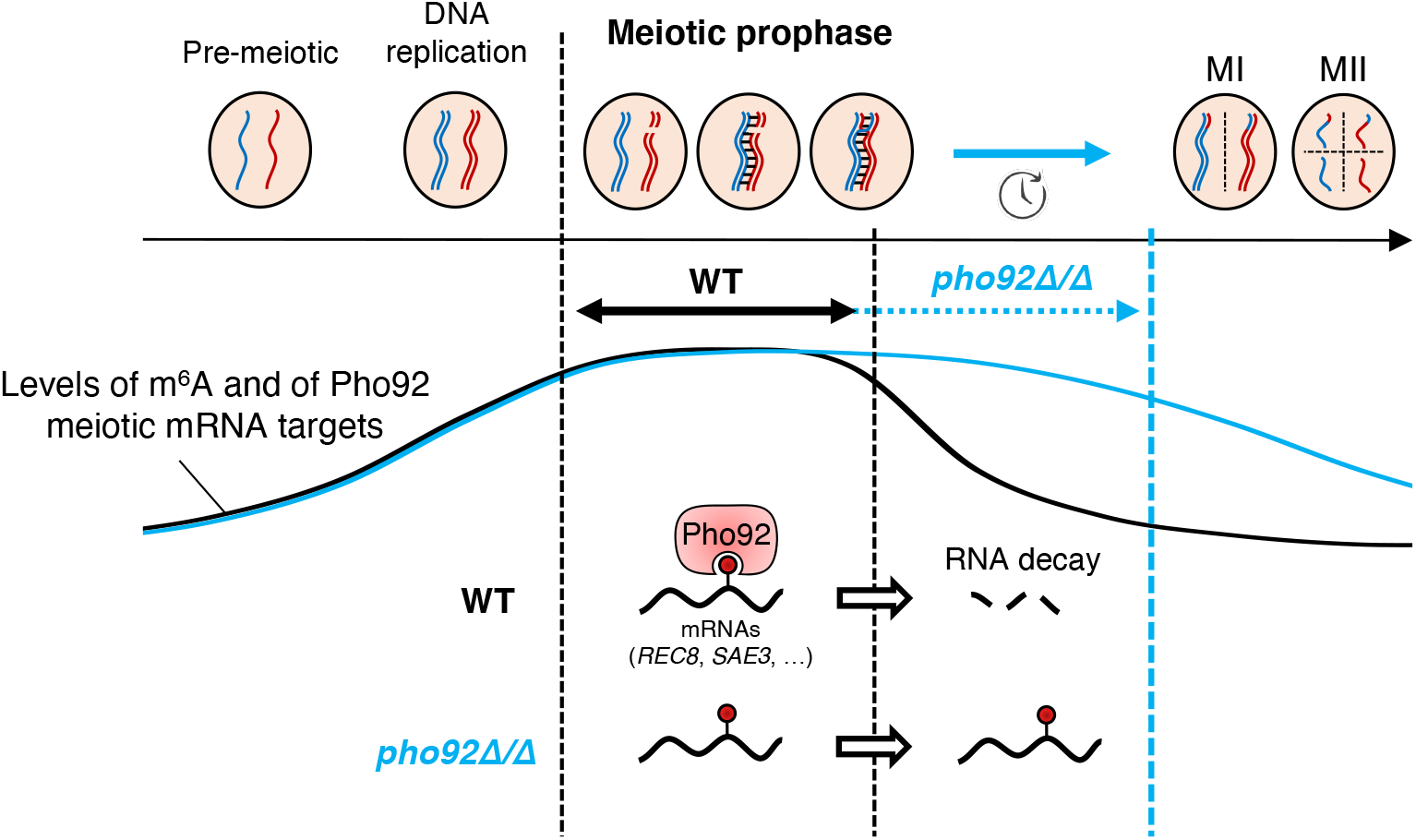
A model for the function of the m^6^A reader Pho92 in the yeast meiotic program. Pho92 promotes the down-regulation of key methylated targets (*REC8* or *SAE3* for example) at late meiotic prophase. The subsequent down-regulation of these mRNAs targets, occurring once they reach their maximum of expression at the onset of meiotic recombination, allows the cells to progress into the downstream events of meiotic divisions. In absence of Pho92, these methylated targets persist and the prophase exit is delayed. A similar persistence of individual mRNA(s) occurs when functional m^6^A site(s) that they carry has (have) been mutated to prevent Ime4-dependent methylation.

Finally, the identification of m^6^A sites bound by the Pho92 reader protein raised the question of whether those were functional. To test this possibility, we mutated sites present in the *SAE3* and *REC8* mRNAs (without altering their coding capacity). Interestingly, these subtle changes were sufficient to impair the down-regulations of these transcripts at late meiotic prophase, similarly to those observed in the absence of Pho92. In the case of *REC8*, the point mutant also impacted the kinetics of meiosis. The latter is consistent with the Rec8 factor being one key actor in temporal control of prophase exit, thus its extended expression potentially impairs the disassembly of the synaptonemal complexes and extends prophase I. In contrast, mutations of the m^6^A site in the *SAE3* mRNA had no detectable consequences on meiosis kinetics. This suggests that the global effect of the absence of Pho92 on meiosis will be the cumulative result of the absence of m^6^A in all of its targets, some possibly speeding the meiotic progression, others having no effect and some delaying it. Thus, our data provide new insights in the complex role of an m^6^A reader in a highly dynamic differentiation program in yeast.

## ACKNOWLEDGMENTS

We acknowledge our team members for support and suggestions. We also thank N. Hollingsworth for the kind gift of antibodies, F. van Werven for kindly sharing the pCUP-IME1 strain, A. Nicolas for the SK1 strain, IGBMC platforms particularly the GenomEast, Imaging, Mediaprep and FACS facilities for support, the I2BC sequencing platform, and D. Heintz and C. Villette for help with m^6^A quantification. The GenomEast platform is a member of the ‘France Génomique’ consortium (ANR-10-INBS-0009) This work was supported by the Ligue Contre le Cancer (Equipe Labellisée 2020) [to B.S.], the Agence Nationale pour la Recherche grant ANR-16-CE11-0003 [to B.S.], the CERBM-IGBMC [to B.S.]. This work of the Interdisciplinary Thematic Institute IMCBio, as part of the ITI 2021-2028 program of the University of Strasbourg, CNRS and Inserm, was supported by IdEx Unistra (ANR-10-IDEX-0002), and by SFRI-STRAT’US project (ANR 20-SFRI-0012) and EUR IMCBio (ANR-17-EURE-0023) under the framework of the French Investments for the Future Program.

## AUTHORS CONTRIBUTIONS

Conceptualization, J.S. and B.S.; Methodology, J.S., M.M. and B.S.; Investigation, J.S., D.P., M.M. T.V., J.Z.; Writing – Original Draft, J.S. and B.S.; Writing – Review & Editing, J.S., D.P., T.V., J.Z., D.L., B.S.; Funding Acquisition, D.L. and B.S.; Supervision, D.L. and B.S.

## EXPERIMENTAL PROCEDURES

### Yeast strains

All yeast strains used in this study are listed in Table S1 and were derived from the sporulation-proficient SK1 background either with an endogenous *IME1* promoter or with *CUP1* inducible promoter driving HA-tagged *IME1* expression (*pCUP*-*IME1*) (Chia and van Werven, 2016). Gene deletions or tagging-fusions were generated in haploid cells with a one-step PCR replacement protocol using standard lithium acetate transformation protocol (Guthrie and Fink, 2002). Pho92 was C-terminally tagged with TAP (CBP-T7-TEV-ProtA) or HTP (His6-TEV-ProtA) using tagging cassette in pBS2438 and pBS6096 (Granneman et al., 2009) respectively (Table S2). Point mutations for *REC8* (A+1461>G) and *SAE3* (C+359>T and A+376>C) were introduced using a “pop-in/pop-out” strategy (Guthrie and Fink, 2002) using the *URA3* marker and 5-FOA counter selection. The presence of desired mutations was determined by PCR and enzymatic digestion, thanks to a polymorphic restriction site introduced in the mutated version (*REC8* point mutant: new BaeGI site; *SAE3* point mutant: new HaeIII site), followed by sequencing of the PCR products. Diploids were obtained by transformation of haploid cells with a YCp50 plasmid carrying the *HO* endonuclease gene with *URA3* selection (pBS433). The presence of the *MAT* a and the *MAT α* genes was verified by PCR on isolated colonies. The *HO* containing-plasmid was lost in rich media before further strain use. Plasmids and oligonucleotides used in this study are listed in Table S2 and Table S3, respectively.

### Growth conditions and meiosis induction

Cells in vegetative growth were grown in YPDA (1% yeast extract, 2% peptone, 2% glucose supplemented with 10 mg/l adenine hemisulfate salt) at 30°C and 170 rpm. Synchronous meiotic entry was performed in standard procedure as described (Schwartz et al., 2013). Briefly, respiratory proficient diploids were pre-selected on 1% yeast extract, 2% peptone, 3% glycerol, 2% agar for 24 hr at 30°C, transferred for another 24 hr of growth in 1% yeast extract, 2% peptone, 4% dextrose at 30°C and 170 rpm, diluted in BYTA media (1% yeast extract, 2% tryptone, 1% potassium acetate, 50 nM potassium phthalate) to OD600 = 0.2 and incubated for another 16 hr incubation at 30°C and 170 rpm. Next, cells were washed once with sterile milliQ water and resuspended in SPO medium (0.3% potassium acetate) at OD600 = 2.0 and incubated at 30°C and 170 rpm. For pCUP-IME1 background, 50 μM of copper (II) Sulfate was added after 2 hr in SPO. Cells were isolated from SPO at the indicated time points by centrifugation (3 min at 8,000 g).

### Western blotting and antibodies

Cells (2 OD600 of meiotic cells or of cells grown in YPDA) were collected by centrifugation and stored at −80°C before use. Total proteins were extracted using the method of Kushnirov (2000). For Figure 4B, proteins were extracted with an improved version of this method (Zhang et al., 2011). Proteins samples were separated on SDS-PAGE (Tris-HCl gels with 8, 10 or 15 % 29:1 acryl:bis-acrylamide) and transferred on Amersham™ Protran™ 0.45 μm nitrocellulose membranes (GE Healthcare, Cat# 10600033) at 4°C for 1 hr at 110V. After Ponceau staining, membranes were blocked in 5% milk PBS-T (1X PBS buffer with 0.1% Tween-20) and incubated overnight at 4°C with one of the following antibodies: anti-HA.11 (1:2,000; Eurogentec, Cat# MMS-101P-500), anti-Stm1 (1:10,000; kind gift from F. Wyers), anti-Hop1 (1:10,000; kind gift from N. Hollingsworth; De Los Santos and Hollingsworth, 1999), anti-Hed1 (1:20,000; kind gift from N. Hollingsworth; Busygina et al., 2008) or anti-Hed1 p-T40 (1:50,000; kind gift from N. Hollingsworth; Callender et al., 2016). After 4 washes in PBS-T, membranes were incubated 1 hr at RT with Peroxidase-AffiniPure Goat Anti-Mouse (1:5,000; Jackson ImmunoResearch, Cat# 115-035-003) or Anti-rabbit IgG, HRP-linked antibody (1:5,000; Cell Signalling, Cat# 7074S). TAP- and HTP-tagged proteins were detected after a unique incubation of the membranes with Peroxidase Anti-Peroxidase Complex produced in rabbit (1:2,000; Sigma, Cat# P1291). Chemiluminescence signals were revealed on an Amersham™ Imager 600 (GE Healthcare) apparatus with Immobilon® Crescendo Western HRP Substrate (Millipore, Cat# WBLUR0500) according to the manufacturers’ instructions.

### Sequences alignment of Pho92 promoter

To align the *PHO92* promoter sequences, the *PHO92* coding and flanking sequences from representative *Saccharomycetaceae* were recovered from the NCBI database using keyword and BLAST searches (Altschul et al., 1990). Sequences upstream of *PHO92*, often extending to the neighboring *WIP1* gene, were aligned using CLUSTAL X (Thompson et al., 1997) and alignments manually edited. Colored alignments were exported from Jalview (Waterhouse et al., 2009).

### Meiotic progression with DAPI counting

Cells (0.3 - 0.5 OD of 2 OD/ml) were isolated from SPO at the indicated times, centrifuged 3 min at 8,000 g, resuspended in 1 volume of 70% ethanol and stored overnight at 4°C. Cells were then pelleted and resuspended in 2 μg/ml DAPI solution in water. Monitoring of meiotic divisions was assayed by counting the number of DAPI masses per cell, in at least 200 cells for each time point, with a Leica DM 4000B microscope equipped with a Hamamatsu ORCA-Flash 4.0 LT C11440 camera.

### RNA extraction, northern blotting and RT-qPCR

Total RNAs were extracted on 2.0 - 3.0 OD of 2 OD/ml yeast cells using a standard hot acid phenol protocol. Frozen cell pellets were resuspended in equal volumes of: 65°C pre-heated “stabilized phenol water-saturated” pH 4.0 (Eurobio, Cat# GEXPHE01), 50 mM sodium acetate pH 5.5 and acid-washed glass beads (0.5 mm; BioSpec Products, Cat# 11079105) and 1/10e volume of 10% SDS. Samples were then incubated at 65°C for 1 min and vortexed for 30 sec, for a total of 3 times, before centrifugation for 10 min (8,000 g at 4°C). Recovered supernatants were re-extracted twice with phenol:chloroform:isoamyl alcohol 25:24:1 (Sigma, Cat# 77617) at 4°C and ethanol precipitated.

For northern blot analysis, 5 μg of total RNAs were fractionated by formaldehyde-agarose gel electrophoresis and transferred to Amersham™ Hybond-XL nitrocellulose membrane (GE Healthcare, Cat# RPN203S) by capillarity. After two rounds of UV auto-crosslink (120,000 μJ/cm2) on UV Stratalinker® 2400 (Stratagene) apparatus, the membranes were probed in hybridization buffer (6X SSC, 2X Denhardt’s reagent, 0.1% SDS) at 60°C with primers (Table S3) radiolabeled at their 5’-end using T4 Polynucleotide Kinase (NEB, Cat# M0201S) and γ-^32^P ATP according to the manufacturer’s instructions. Radioactive signals were measured using a Typhoon FLA 9000 Phosphorimager (GE Healthcare).

For quantitative RT-PCR, total RNA samples were treated with DNase I recombinant, RNAse-free (Roche, Cat# 4716728001) at 37°C for 1 hr and purified by phenol-chloroform extraction as described before. cDNAs were generated from 500 ng of DNase-treated RNAs with SuperScript™ IV reverse transcriptase (Invitrogen™, Cat# 18090050) and oligo(dT)18 primer (Thermo Scientific™, Cat# S0132) according to the manufacturers’ instructions. Quantitative PCRs were performed using LightCycler® SYBR Green I Master (Roche, Cat# 04887352001) with primers listed in Table S3 on a LightCycler® LC480 instrument (Roche). Primer efficiencies were > 95% and calculated on standard curves generated by a 2-fold dilution series of cDNAs over at least five dilution points measured in triplicate. Each sample was measured in biological triplicates with technical duplicates and transcript abundances were determined using the 2^−ΔΔCt^ method normalized to *ACT1*.

### m^6^A quantification by mass spectrometry

Total RNAs were extracted from yeast meiotic cells as described before and polyadenylated RNAs were isolated from 50 μg total RNAs as starting material with two successive rounds of poly-A selection using Dynabeads™ mRNA Purification kit (Invitrogen™, Cat# 61006) according to the manufacturer’s instructions. Absence of rRNA contamination and mRNA integrity were verified with Agilent Bioanalyzer. One hundred nanograms of isolated poly(A)+ RNAs was digested into single nucleosides with 1 U of benzonase® nuclease (Millipore, Cat# E1014-5KU), 0.003 U of Phosphodiesterase I (Sigma, Cat# P3243-1VL) and 1 U of shrimp alkaline phosphatase (NEB, Cat# M0371S) in a 50 μL buffer containing 10 mM Tris-HCl pH 8.0, 1 mM MgCl2 and 0.1 mg/ml BSA. After an incubation for 6 hr at 37°C, samples were filtered on Nanosep 3K omega tubes (Pall, Cat# OD003C34) by centrifugation at 12,000 g for 10 min. Ten microliters aliquots were analyzed by LC-MS/MS analysis on an EvoQ Elite TQ (Bruker) equipped with a Dionex UltiMate 3000 UHPLC system (Thermo) coupled to an electrospray ionization source (ESI). Chromatographic separation was achieved using an Acquity UPLC HSS T3 column (100 x 2.1 mm, 1.8 µm; Waters) and pre-column (5×2.1mm, 1.8µm; Waters). The mobile phase consisted of (A) water and (B) methanol, both containing 0.1 % formic acid. The run started by 0 min of 98 % A, then a convex gradient (curve number 5) was applied to reach 7 % B at 4 min then another 8-min concave gradient (curve number 8) to reach 100% B and maintained 0.5 min. Return to initial conditions was achieved in 1 min and maintained 1.5 min. The total run time was 15 min. The column was operated at 35 °C with a flow-rate of 0.32 mL/min. Nitrogen was used as the drying and nebulizing gas. The nebulizer gas flow was set to 35 L/h, and the desolvation gas flow to 30 L/h. The interface temperature was set to 350 °C and the source temperature to 300 °C. The capillary voltage was set to 3.5 kV; the ionization was in positive mode. Low mass and high mass resolution were 2 for the first mass analyzer and 2 for the second. The transitions were, in positive mode: adenosine 268.0>136.10 and 268.0>119.10 (collision energy 15V and 42V respectively); N^6^-methyladenosine 281.9>150.10 and 281.9>123.10 (collision energy 17V and 41V respectively). Data acquisition was performed with the MS Workstation 8 for the mass spectrometry and the liquid chromatography was piloted with Bruker Compass Hystar 4.1 SR1 software and Chromeleon Xpress (Thermo). The data analysis was performed with the Bruker MS Data Review software (ver 8.2.1).

Standard solutions of adenosine (0.05, 0.1, 0.5, 1, 5, 10 ng/ml; Sigma, Cat# A9251) and N^6^-methyladenosine (0.05, 0.1, 0.5, 1, 5 ng/ml; Selleckchem, Cat# S3190) were used for the quantification. All the calculated values for the different nucleosides in each sample fell within the standard curve range. The ratio of N^6^-methyladenosine to adenosine was calculated based on the calibrated concentrations.

### Transcriptome analyses

mRNA-seq libraries were generated from 500 ng of total RNA using TruSeq® Stranded mRNA Library Prep kit and TruSeq® RNA Single Indexes kits A and B (Illumina, Cat#20020595) according to manufacturer’s instructions. Quantity and quality of the started material and final librairies were checked using a Qubit™ fluorometer (Invitrogen™) and Agilent Bioanalyzer. Resulting libraries were sequenced in 50-length Single-Read on a HiSeq 4000 machine (Illumina).

#### Reads mapping

Reads were mapped onto the SK1 assembly of the *Saccharomyces cerevisiae* genome (SK1 MvO V1; SK1_SGD_2018_NCSL00000000) using Tophat 2.0.14 (Kim et al., 2013; Trapnell et al., 2009) and bowtie version 2-2.1.0 (Langmead et al., 2009). Only uniquely mapped reads have been retained for further analyses. Quantification of gene expression has been performed with HTSeq-0.6.1 (Anders et al., 2015), using intersection-nonempty mode, and annotations coming from the Saccharomyces Genome Database (SGD). Only non-ambiguously assigned reads have been retained for further analyses.

#### Clustering

Genes with larger average expression have on average larger observed variances across samples, that is, they vary in expression from sample to sample more than other genes with lower average expression. To avoid these genes to overly influenced the variance, variance have been stabilized using the rlog (Regularized log) function from DESeq2. Principal Component Analysis (PCA) is based on their normalized and stabilized gene expression level of all samples.

#### Statistical analysis

Comparisons of interest have been performed using the R 3.5.1 with *DEseq2* version 1.22.1 (Love et al., 2014). More precisely, read counts were normalized from the estimated size factors using the median-of-ratios method (Anders and Huber, 2010) and a Wald test was used to estimate the p-values. P-values were then adjusted for multiple testing with the Benjamini and Hochberg method (Benjamini and Hochberg, 1995).

#### Temporal clustering

In order to have a better view of the changes in expression of significant genes between each time, a clustering using the Mfuzz (2.42.0) R package (Kumar and Futschik, 2007) has been realized. Only genes with an adjusted p-value lower than 0.05 in the WT time-series analysis and having a | log2 FoldChange | > 2 (calculated according to the time point having the maximum count relative to the time point having the minimum count) has been retained for clustering. To measure the impact of Pho92 and Ime4 deletion, the expression value of the corresponding genes in the mutants were plotted according to the clustering established for the WT strain. Heat maps were build using “heatmapper” online tool (Babicki et al., 2016).

### Flow cytometry analysis of meiotic DNA replication

Two hundred microliters of meiotic cells (at 2 OD/ml) were collected at each time point, resuspended in 70% ethanol and stored overnight at 4°C. Samples were then spun down, resuspended in 200 μL of solution A (50 mM sodium citrate, 0.1 mg/ml RNAse A (Sigma, Cat# R6513), 0.2 mg/ml Proteinase K (Thermo Scientific™, Cat# EO0491)) and incubated 2 hr at 50°C. Cells were pelleted again and resuspended in 500 μl of 50 mM sodium citrate containing 2 μM of SYTOX Green (Invitrogen™, Cat# S7020) and transferred to 5 ml FACS tubes (Becton Dickinson, Cat# 352008). After vortexing, samples were analyzed on the LSR II (BD Biosciences) instrument and BD FACS Diva software. Single cells were gated based on forward and side scattering, FITC-A, and FITC-W. Ten thousand events were counted per sample. Data were analyzed using FlowJo 10 software.

### Detection of meiotic double strand breaks by southern blotting

Yeast cells were collected every hour after transfer to SPO (at 2 OD/ml) as above and genomic DNA was extracted on equivalent of 30 OD (time points: 0 hr, 2 hr, 3 hr after transfer to SPO) or 15 OD (time points: 4 hr to 9 hr) according to the method described for medium-resolution mapping of double strand breaks (Murakami et al., 2009). Up to 5 μg of gDNA was digested with 15 U of SalI-HF® enzyme (NEB, Cat# R3138L) for 3 hr at 37°C and separated on 0.9% agarose gel for 16 hr at 50 V. Next, the gel was incubated 45 min with shaking in 0.2 N NaOH, 0.6 M NaCl and 45 min in 1 M Tris-HCl pH 7.5, 0.6 M NaCl before transfer under neutral conditions on Amersham™ Hybond-XL nitrocellulose membrane (GE Healthcare, Cat# RPN203S). After one round of UV auto-crosslink (120,000 μJ/cm2) on UV Stratalinker® 2400 (Stratagene) apparatus, the membrane was hybridized with a random-primed 32P-labeled specific probe using Amersham™ Megaprime kit (GE Healthcare, Cat# RPN1607). The probe was obtained from a 1.45 kb PCR product (primers list in Table S3) covering a region of the*YCR048W* gene.

### Luc7-HTP tagging in *S. pombe*

*Schizosaccharomyces pombe* cells (972 h-background) were grown at 30°C in YES media (1% yeast extract, 3% glucose, 225 mg/l adenine, uracil, leucine and lysine). C-terminal HTP-tagging of the essential U1-snRNP associated protein Luc7 (SPCC16A11.13) was obtained with a one-step PCR replacement method using a pFA6a-HTP-KanMX6 (pBS6122) plasmid template and introduced with a standard lithium acetate transformation protocol (Rai et al., 2018). Positive transformants were selected on YES + G418 (200 μg/ml) and verified by PCR. Luc7-HTP functional expression was validated by western blot on total protein extracts (Matsuo et al., 2006) obtained from 0.5 OD of cells.

### Crosslinking and analysis of cDNA (CRAC)

CRAC was performed essentially as described (Villa et al., 2020) with minor modifications. To facilitate later normalization between samples during dataset analysis, we used an internal “spike-in” extract of *S. pombe* expressing an RNA-binding protein fused to the HTP tag. The essential protein Luc7 (30.94 ± 19.0 kDA HTP) with a comparable molecular weight to *S. cerevisiae* Pho92 (36.05 ± 19.0 kDA HTP) was chosen in order to collect both proteins in the same fraction during the gel-free protein fractionation step of the CRAC protocol (see below).

Meiosis was induced as previously described in 500 mL of SPO to OD600 = 2.0 at 30°C and 5 hr following resuspension in SPO (corresponding to 3 hr after *IME1* induction), samples were diluted to a final volume of 2 L with SPO media and UV-irradiated at 254 nm for 80 s using the Megatron W5 UV crosslinking unit (UVO3 Ltd). The control samples “Pho92-HTP No UV” were directly processed without UV-irradiation. The sample used as “spike-in” from *S. pombe* cells expressing Luc7-HTP was grown independently in 800 mL of 1X EMM media (EMM Broth, Formedium, Cat #PMD0210) supplemented with 2% glucose and 125 mg/L each of adenine (Sigma, Cat #A2786), L-histidine (Sigma, Cat #H8000), uracil (Sigma, Cat #U750), L-lysine (Sigma, Cat #L5501) and L-leucine (Sigma, Cat #L8000). Cells were grown to 0D600 = 0.5 at 30°C and then diluted to a final volume of 2 L before UV-irradiation (254 nm for 80 s). All samples were harvested by centrifugation, washed in 35 ml cold PBS and resuspended in 2.5 ml per g of cells (or 1.25 ml/g for *S. pombe* cells) of TN150 buffer (50 mM Tris pH 7.8, 150 mM NaCl, 0.1% NP-40 and 5 mM 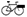-mercaptoethanol) supplemented with 0.4 μM AEBSF (Sigma, Cat# A8456), 0.4 μM Benzamidine (Sigma, Cat# B6506 and cOmplete™, EDTA-free Protease Inhibitor Cocktail (Roche, Cat# 11873580001). The suspension was flash frozen in droplets, then mechanically broken through 5 cycles of 3 minutes at 15 Hz in a Mixer Mill MM 400 (Retsch, Cat# 20.745.0001) and resulting cell powders stored at −80°C until use.

Cell powders were thawed and an amount of *S. pombe* lysate corresponding to 50 OD600 (roughly 0.2 g of cells) was added to each *S. cerevisiae* sample, giving a final 5% fraction of *S. pombe* over *S. cerevisiae* cells based on OD600. The resulting lysates were treated for 1 hr at 18°C with DNase I (165 U/g of cells) in the presence of 10 mM MnCl_2_ to solubilize chromatin and then centrifuged at 4,800 g for 20 min at 4°C. The supernatant was moved to a fresh tube and further clarified by centrifugation at 22,000 g for 30 min at 4°C. Cleared lysates were incubated with 200 μL Dynabeads™M-280 tosylactivated (Invitrogen™, Cat#14203) coupled with rabbit IgG (15 mg of beads per samples; Sigma, Cat# I5006), nutating at 4°C for 2 hr. Beads were washed three times with 10 mL TN1000 (same as TN150 + protease inhibitors, but with 1 M NaCl) for 5 min, and once with 10 mL TN150. His-tagged protein-RNA complexes were eluted from IgG beads with 5 μL homemade recombinant GST-TEV protease for 2 hr at 18°C with shaking in 600 μL TN150 supplemented with 0.4 μM oligo (dT), 3 mM MgCl_2_ and 2 μL (10 U) RNase H (NEB, Cat# M0297S) in order to digest poly(A) tails from RNAs at the same time, thus favoring subsequent reads mapping. Eluate were then treated with 0.1 U RNace-IT RNase cocktail (Agilent, Cat# 400720) for 5 min at 37°C to fragment protein-bound RNA. The RNase reaction was quenched with the addition to 400 mg guanidine hydrochloride. The solution was adjusted for nickel affinity purification to 0.3 M NaCl and 15 mM imidazole, added to 80 μl washed Ni-NTA agarose nickel beads slurry (Qiagen, Cat# 30230), and transferred to Pierce™ Spin Columns – Snap Cap (Thermo Scientific™, Cat# 69725).

Following an overnight incubation at 4°C, nickel beads were washed twice with Wash Buffer I (6.0 M guanidine hydrochloride, 50 mM Tris-HCl pH 7.8, 300 mM NaCl, 0.1% NP-40, 10 mM imidazole, and 5 mM *β*-mercaptoethanol), and then three times with 1x PNK buffer (50 mM Tris-HCl pH 7.8, 10 mM MgCl_2_, 0.1% NP-40, and 5 mM *β*-mercaptoethanol). Subsequent reactions were performed on the columns in a total volume of 80 μl, ending each time by one wash with Wash Buffer I and three washes with 1x PNK buffer, in the following order:

1. Phosphatase treatment (1x PNK buffer, 50 U QuickCIP (NEB, Cat# M0525L), 80 U RNaseOUT (Invitrogen, Cat#10777019); 37°C for 30 min).
2. 3’ linker ligation (1x PNK buffer, 800 U T4 RNA ligase 2 truncated KQ (NEB, Cat#M0373L), 80 U RNaseOUT, 1 μM preadenylated 3’ linkers, modified for sequencing from the 3’ end (list in Table S3); 25°C for 5.5 hr).
3. 5’ end phosphorylation (1x PNK buffer, 20 U T4 PNK (NEB, Cat# M0201L), 80 U RNaseOUT, 1.5 mM ATP; 37°C for 45 min).
4. 5’ linker ligation (1x PNK buffer, 40 U T4 RNA ligase I (NEB, Cat# M0204L), 80 U RNaseOUT, 1.25 μM 5’ linker (L5 miRCat), 1 mM ATP; 16°C overnight).

The beads were washed three times with Washing Buffer II (50 mM Tris-HCl pH 7.8, 50 mM NaCl, 0.1% NP-40, 10 mM imidazole, and 5 mM *β*-mercaptoethanol). Protein-RNA complexes were eluted for 2 x 10 min in 200 μl elution buffer (same as Washing Buffer II but with 150 mM imidazole). Eluates from control samples (Pho92-HTP No UV x 2 rep + WT x 2 rep) were mixed together as well as eluates from samples (Pho92-HTP x 2 reps + Pho92-HTP *ime4Δ/Δ* x 2 reps) before concentration with Vivacon® 500 filtration cartridges (10 kDa MWCO; Sartorius, Cat# VN01H02) to a final volume of 120 μL. The protein fractionation step was performed with a Gel Elution Liquid Fraction Entrapment Electrophoresis (Gelfree 8100) system (Expedeon, Cat# 48100). Final fractions were treated with 100 μg of proteinase K (Roche) in buffer containing 0.5 % SDS for 2 hr at 55°C with shaking. RNA was isolated with phenol:chloroform extraction followed by ethanol precipitation.

RNA was reverse transcribed using SuperScript™ IV reverse transcriptase (Invitrogen™, Cat# 18090050) and the RT L3-2 oligo for 1 hr at 50°C in a 20 μL reaction. Samples were heat inactivated (80°C, 10 min) and then treated with 1 μL (5 U) RNase H (37°C, 30 min). The absolute concentration of cDNAs in the reaction was estimated by quantitative PCR using a standard of known concentration. Then, cDNA was amplified by PCR in separate 25 μL reactions each containing 2 μL of cDNA for 11-13 cycles using 0.5 μL (2.5 U) LA Taq (Takara, Cat# RR002M) with P5-3’ and miRCat-PCR-2 oligos at an annealing temperature of 58°C. The PCR reactions were pooled and treated for 1 hr at 37°C with 25 U of Exonuclease I (NEB, Cat# M0293S) per 100 μL of pooled PCR reactions. Libraries were purified using NucleoSpin Gel and PCR Clean-up (Macherey-Nagel) and quantified with a Qubit Fluorometer and the Qubit dsDNA HS Assay Kit (Invitrogen, Cat# 740609). Single-end sequencing was performed using Illumina technology on a NextSeq 500 instrument.

### CRAC peak calling

CRAC datasets were analyzed as previously described (Candelli et al., 2018) except for the normalization step. Briefly, data preprocessing steps include the 5’ adapter trimming, PCR duplicates and poly-A stretch removal and the reverse complement of the reads. BAM files were generated using bowtie2 (2.4.1) and the *Saccharomyces cerevisiae* genome (SK1 assembly). The peaks were detected using the extractPeaks pipeline of the PeakCcall suite after gaussian kernel smoothing of the CRAC signal [Challal et al., MolCell 2018] and scored according to the “h^2^w” score, where “h” = height, and “w” = width of the peak. The normalization between samples was done according to the internal Luc7-HTP spike-in. For that, the reads that did not map to the S.cerevisiae SK1 genome were mapped to the *Schizosaccharomyces pombe* genome (version ASM294v2) after filtering for a minimal length of 40 nt to avoid cross-species mapping. PeakCcall was used to determine the binding score of Luc7-HTP to its U1 snRNA target at the locus (chr II:3020203-3020352) in each sample. The resulted score was used to normalize the score of all *S. cerevisiae* peaks of the respective sample and relative to “Pho92-HTP Replicate 1” sample. Only the peaks having a score > 0.9 were considered as *bona fide* peaks, corresponding to scores above the 98.4 percentiles of the distribution of all peak scores in the “Pho92-HTP Replicate 1” sample. The intersection of the peaks was made between the two replicates of Pho92-HTP according to the barycentre coordinate of each peak for which −3 and +3 bases were added, using intersectBed from BEDTools (2.21.0). Annotation of the peaks was carried out using closestBed from BEDTools and the annotation file converted in BED format.

### Motif discovery, metagene analysis and Gene Ontology

*De novo* motif discovery was performed with the MEME suite (5.4.1) (Bailey et al., 2015) with the default parameters except for the motifs size fixed to 9 nucleotides. Fasta sequences corresponding to the different populations of extracted CRAC peaks were used as input.

UTR extensions have been defined according to the file generated in Bushkin et al. (2019). The mean length of the 5’UTR, CDS and 3’UTR were calculated and the position of each peak relative to the corresponding mean was determined. To put together the positions of each part of the transcript into a figure, the mean length of the 5’UTR was added to the peak position on the CDS and the mean length of the 5’UTR and the CDS were added to the peak position on the 3’UTR.

Gene ontology was performed either: for Figure 3C, with the GO Term Finder tool from the *Saccharomyces* Genome Database (yeastgenome.org/goTermFinder) using default parameters and a p-value threshold of 0.05; or for Figure 5E, with ShinyGO v0.75 (Xijin Ge et al., 2020) using a P-value cutoff (FDR) of 0.05.

### MazF-qPCR assay for determination of methylation sites

The MazF digestion and qPCR design were based on a described method (Garcia-Campos et al., 2019) with few modifications. Briefly, 500 ng of total RNAs, isolated as described before at 3 and 7 hr after transfer in SPO, were incubated 2 min at 70°C and placed on ice. Each sample was then incubated at 37°C for 30 min with 20 U of mRNA-Interferase™ MazF enzyme (Takara, Cat# 2415A) in a 10 μL volume reaction containing 1X MazF reaction buffer, 40 U of RNasin® Ribonuclease Inhibitors (Promega, Cat# N2511). All samples were duplicated to include control samples without the MazF enzyme. Reactions were stopped by placing on ice and cDNAs were directly generated from half of each sample using the SuperScript™ IV reverse transcriptase (Invitrogen™, Cat# 18090050) and random hexamer primer (Thermo Scientific™, Cat# SO142). qPCRs were performed as described above using two primers pairs per tested gene. The first “target” primer pair was designed to flank a putative methylation site with only one ACA site corresponding to the interrogated site, thus amplified only if methylated and not digested by MazF. The second “control” primer pair was designed to flank an adjacent region in the same gene without an ACA site. The methylation scores of the interrogated site were estimated using the following formula: E (efficiency) = 2^−[(Ct MazF+)-(Ct MazF-)]^ and methylation scores = E (“target”)/E (“control”) (Garcia-Campos et al., 2019). It relates the transcript abundance of the “target” amplicon in MazF-treated versus untreated samples, normalized to the transcript abundance of the “control” amplicon in MazF-treated versus untreated samples.

