## Supplemental Figures S1-S6 for "The *S. cerevisiae* m^6^A-reader Pho92 impacts meiotic recombination by controlling key methylated transcripts"

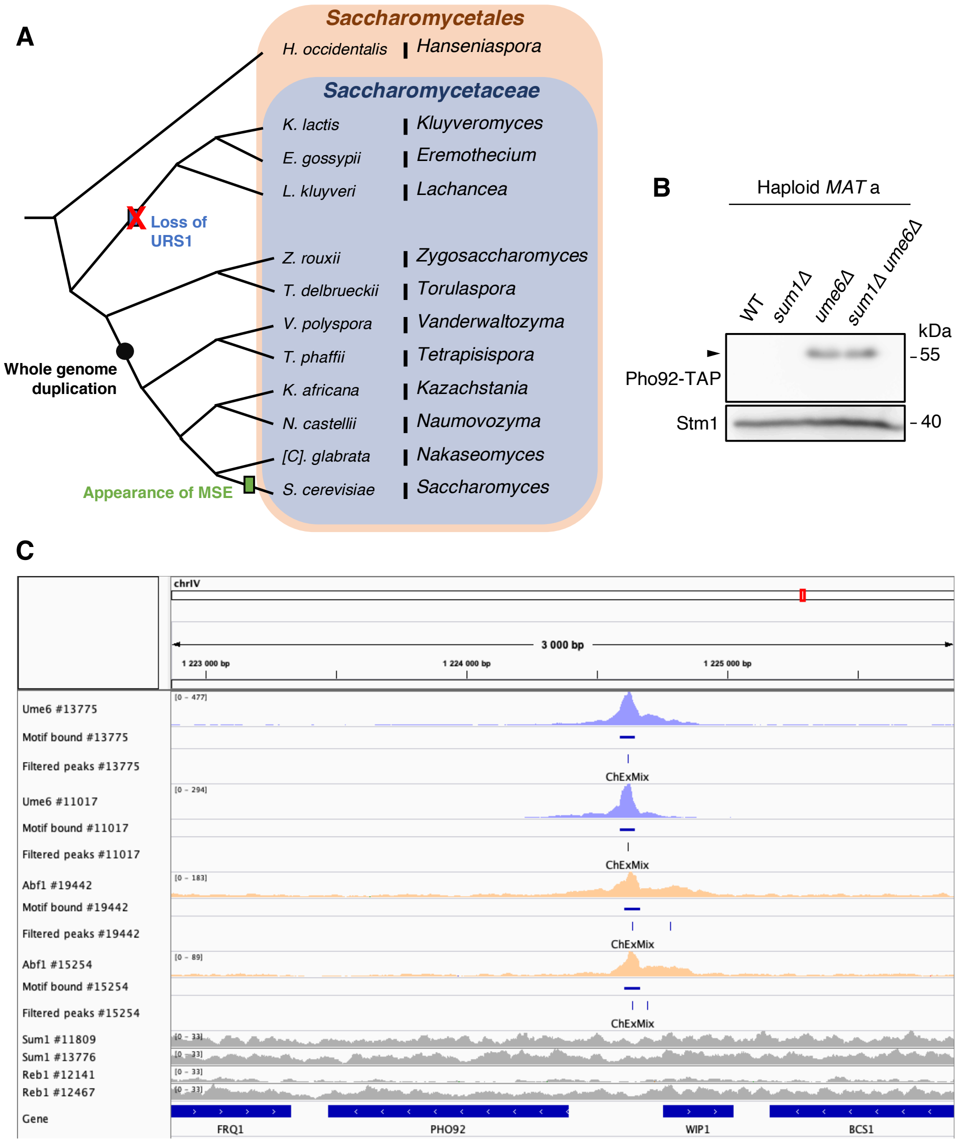


**Figure S1 (related to Figure 1). Elements in the promoter of *PHO92* participate in its transcription regulation.**

(**A**) Tree indicating events in the evolution of the Pho92 promoter in *Saccharomycetaceae*, the *Saccharomycetales Hanseniaspora occidentalis* species being used for rooting. Loss of URS1, whole-genome duplication and appearance of MSE are indicated. Only a subset of species present in Figure 1C are indicated.

(**B**) Expression of Pho92-TAP in haploid strains of the SK1 background grown in rich media (YPDA). Western blot using the strains as in Figure 1D in their haploid versions.

(**C**) Integrative Genome Browser (IGV) tracks of Ume6, Abf1, Reb1 and Sum1 binding data in the genomic region of Pho92. Original data were obtained by ChIP-exo-seq (chromatin immuno-precipitation, exonuclease digestion and DNA sequencing) in haploid cells grown in rich media (Rossi et al., 2021). The tracks represent the read coverage, the region of the motif bound and the position of final extracted peaks for two biological replicates of each transcription factor, according to the #ID in the original data.

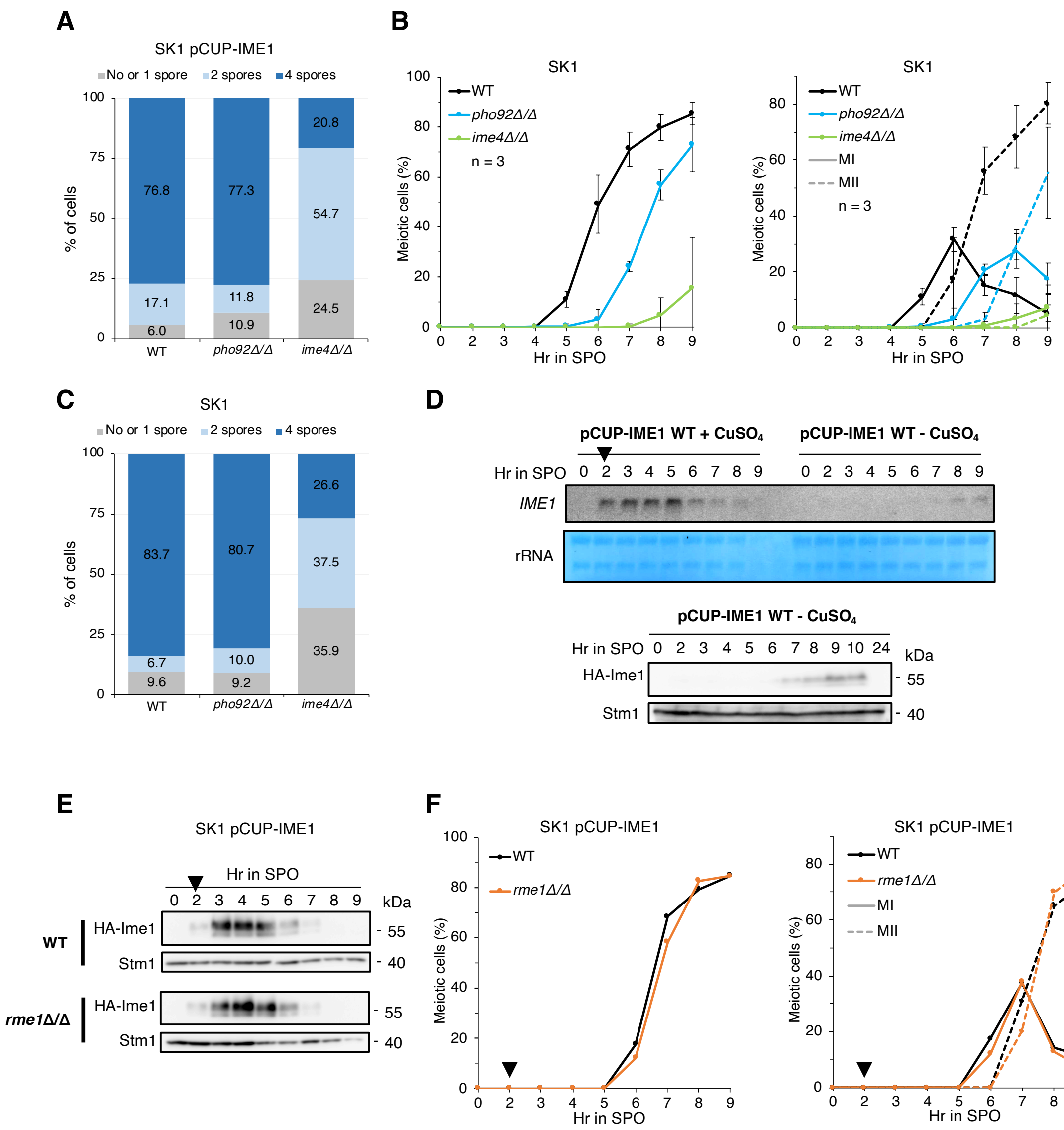


**Figure S2 (related to Figure 2). Meiotic phenotypes in different strain backgrounds, *RME1* has no impact on *IME1* expression and meiosis in pCUP-IME1 background.**

(**A**) Quantification of asci with no/1, 2 or 4 spores after 24 hr in SPO medium from WT, *pho92Δ/Δ* and *ime4Δ/Δ* cells in the pCUP-IME1 background. The number of spores was monitored in 200 cells by counting the number of nuclei per asci after DAPI staining. Mean values from 3 biological replicates.

(**B**) Kinetics of meiotic cells appearance in WT, *pho92Δ/Δ* and *ime4Δ/Δ* cells of the SK1 background following resuspension in SPO medium. The number of nuclei per cell was monitored by microscopic observation of DAPI staining in 200 cells per time point. Left panel: total number of meiotic cells containing >1 nucleus. Right panel: cells completing meiosis I (solid lines) and meiosis II (dashed lines). Means and standard deviations from 3 biological replicates.

(**C**) Same as (A) in strains in the SK1 background. Mean values from 3 biological replicates. (**D**) Analysis of *IME1* expression, using northern blot (upper panel) and western blot (lower panel) with an anti-HA antibody, when no copper (II) sulfate is added after 2 hr in SPO medium in WT cells in the pCUP-IME1 background. Total RNAs from WT cells with induction at 2 hr SPO of *IME1* (black triangle) were used as comparison in the northern blot analysis. Loading controls are rRNA signals from methylene blue staining for the northern blot and Stm1 signal for the western blot. Note that even if low levels of Ime1 protein are detected at late time points, no meiotic cells were detected after 10 hr in SPO.

(**E**) Western blot using anti-HA antibody to detect HA-Ime1 in total protein extracts from WT and *rme1Δ/Δ* mutant cells collected at different time points following resuspension in SPO medium and induction of *IME1* at 2 hr (black triangle). Stm1 signal detected in the same extracts was used as loading control.

(**F**) Kinetics of meiotic divisions in samples collected for (E). The number of nuclei per cell was assayed by DAPI staining in 200 cells per time point. Left panel: total number of meiotic cells containing >1 nucleus. Right panel: cells completing meiosis I (solid lines) and meiosis II (dashed lines). Black triangle above 2 hr: time of *IME1* induction.

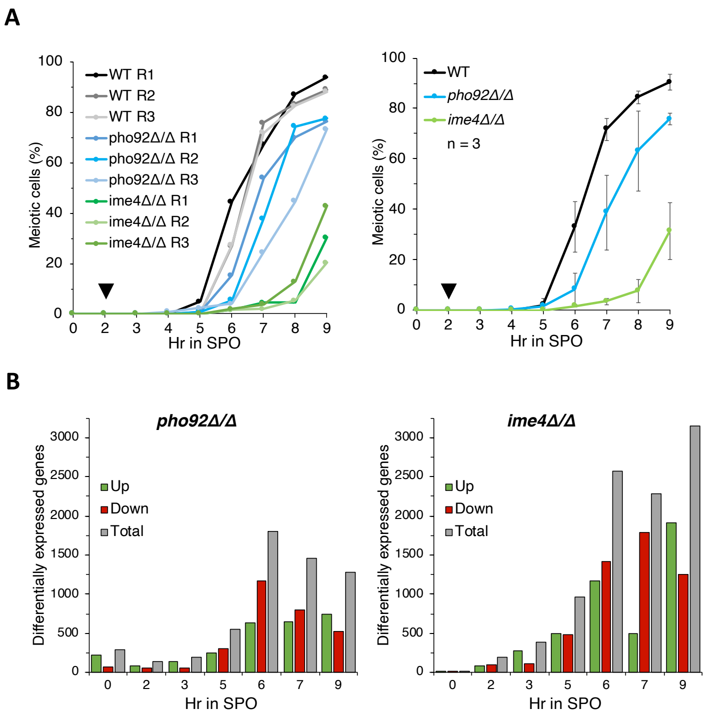


**Figure S3 (related to Figure 3). Meiotic progression of samples used for transcriptomic analyses and number of differentially expressed genes.**

(**A**) Kinetics of meiotic cells appearance in samples collected for mRNA-sequencing. The number of nuclei per cell was assayed by DAPI staining in 200 cells per time point. Left panel: individual values of meiotic cells (>1 nucleus) distribution in three biological replicates (R1, R2, R3) of WT, *pho92Δ/Δ* and *ime4Δ/Δ* cells following resuspension in SPO medium and induction of *IME1* after 2 hr (black triangle) by addition of copper (II) sulfate. Right panel: Means and standard deviations from the 3 biological replicates.

(**B**) Number of differentially expressed genes in *pho92Δ/Δ* (left panel) and *ime4Δ/Δ* (right panel) mutants compared to WT at each time point of the meiotic time course (grey). Only genes presenting a p-value adjusted for multiple testing <0.05 were considered and defined as up-regulated if log2[Fold change(mutant/WT)] >1 (green) or down-regulated if log2[Fold change(mutant/WT)] <-1 (red).

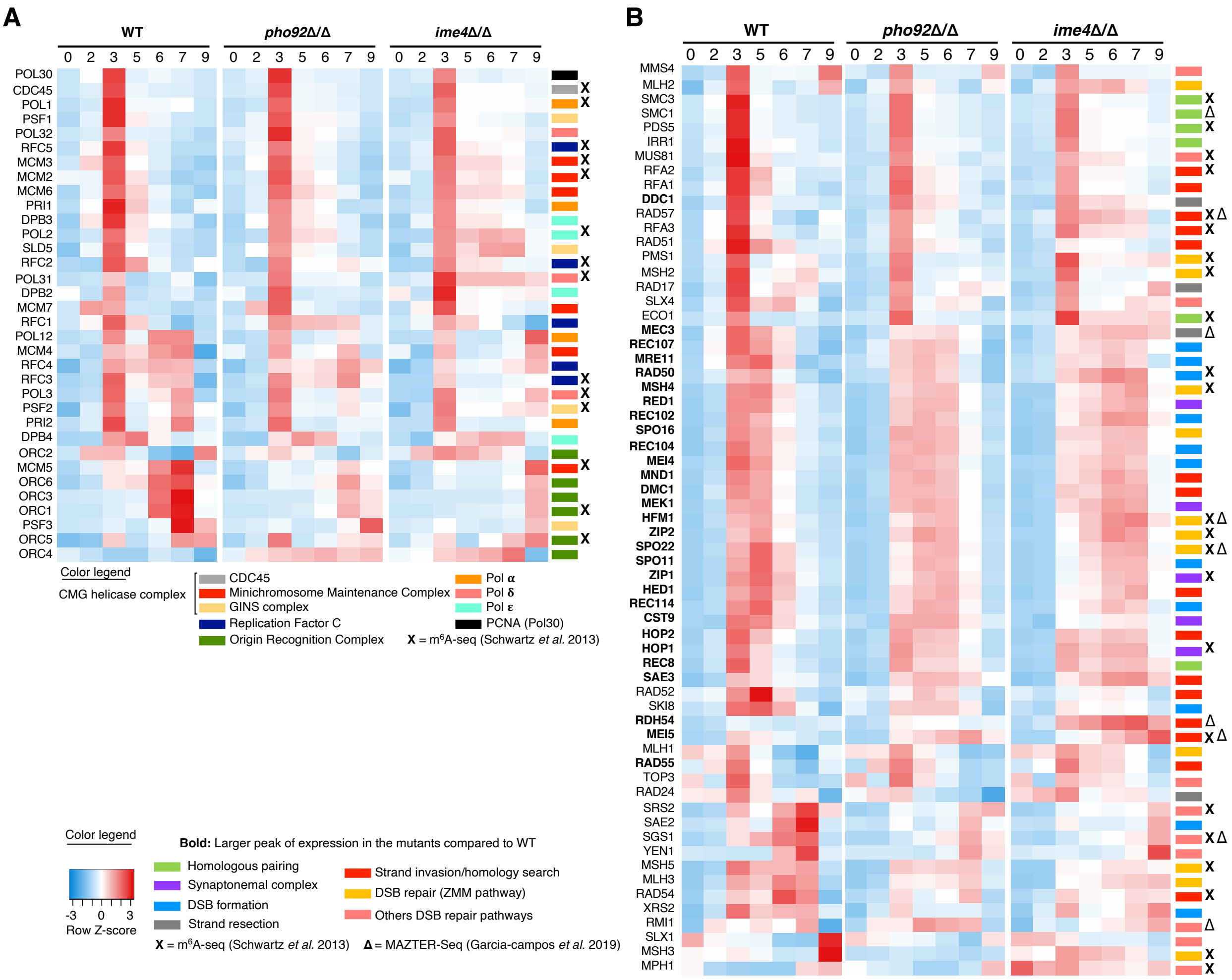


**Figure S4 (related to Figure 4). Pho92 affects the expression of meiotic DSB formation- and processing-related genes but not the induction of DNA replication factors.**

(**A**) Heat map representation of expression of key genes involved in DNA replication following meiosis induction in WT, *pho92Δ/Δ* and *ime4Δ/Δ* cells. Z-score transformation was performed for each gene considering the mean normalized read count of three biological replicates. Each row represents one gene and each column represents a sample. Z-score is scaled for each line from red (maximum expression value from all samples) to blue (lowest expression value from all samples). Genes are arranged together according to the chronology of their respective maximum expression in the WT strain. At right, a supplementary color code indicates the protein complex in which each gene product is involved. Black cross: methylated candidates in m$^6$A-seq data (Schwartz et al., 2013).

(**B**) Same as (A) but considering a list of key genes involved in meiotic recombination. At right, the color code indicates the major meiotic function or step in which each gene product is involved. Genes previously identified as methylated candidates are indicated with a black cross (Schwartz et al., 2013) or black triangle (Garcia-Campos et al., 2019; confidence group >1).

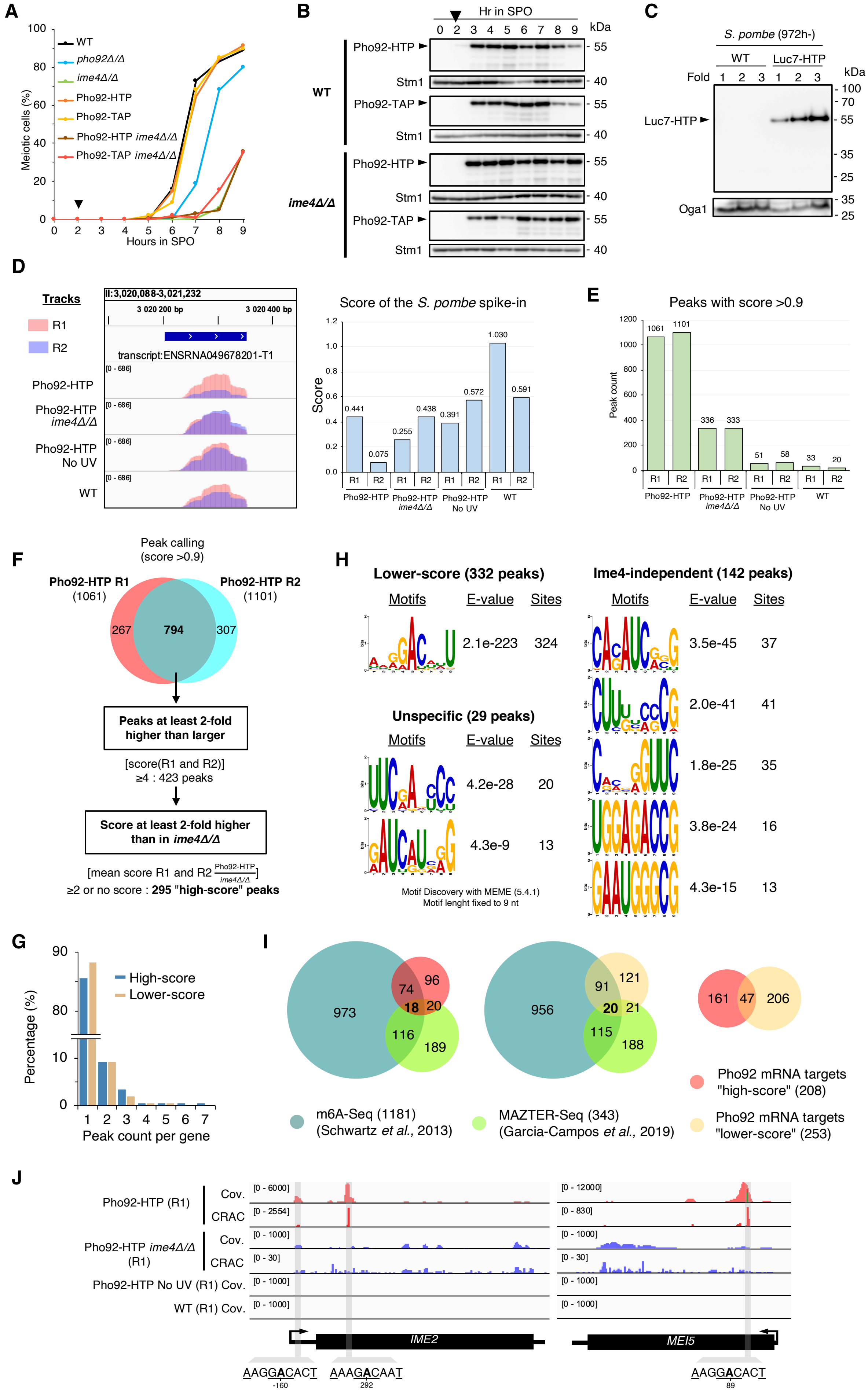
A
Meiotic cells (%)
Hours in SPO
WT
pho92Δ/Δ
ime4Δ/Δ
Pho92-HTP
Pho92-TAP
Pho92-HTP ime4Δ/Δ
Pho92-TAP ime4Δ/Δ
B
Hr in SPO
0 2 3 4 5 6 7 8 9
kDa
WT
ime4Δ/Δ
Pho92-HTP
Stm1
Pho92-TAP
C
S. pombe (972h-)
WT
Luc7-HTP
Fold 1 2 3 1 2 3
Luc7-HTP
Oga1
D
Tracks
R1
R2
II:3,020,088-3,021,232
3 020 200 bp
3 020 400 bp
transcript:ENSRNA049678201-T1
Pho92-HTP
Pho92-HTP ime4Δ/Δ
Pho92-HTP No UV
WT
Score of the S. pombe spike-in
Score
0.441
0.075
0.255
0.438
0.391
0.572
1.030
0.591
E
Peaks with score >0.9
Peak count
1061
1101
336
333
51
58
33
20
F
Peak calling (score >0.9)
Pho92-HTP R1 (1061)
Pho92-HTP R2 (1101)
267
794
307
Peaks at least 2-fold higher than larger
[score(R1 and R2)] ≥4 : 423 peaks
Score at least 2-fold higher than in ime4Δ/Δ
[mean score R1 and R2 Pho92-HTP/ime4Δ/Δ] ≥2 or no score : 295 "high-score" peaks
G
Percentage (%)
High-score
Lower-score
Peak count per gene
H
Lower-score (332 peaks)
Motifs E-value Sites
2.1e-223 324
Unspecific (29 peaks)
Motifs E-value Sites
4.2e-28 20
4.3e-9 13
Motif Discovery with MEME (5.4.1)
Motif lenght fixed to 9 nt
Ime4-independent (142 peaks)
Motifs E-value Sites
3.5e-45 37
2.0e-41 41
1.8e-25 35
3.8e-24 16
4.3e-15 13
I
973
74
96
18
20
116
189
956
91
121
20
21
115
188
161
47
206
m6A-Seq (1181) (Schwartz et al., 2013)
MAZTER-Seq (343) (Garcia-Campos et al., 2019)
Pho92 mRNA targets "high-score" (208)
Pho92 mRNA targets "lower-score" (253)
J
Pho92-HTP (R1)
Cov.
CRAC
Pho92-HTP ime4Δ/Δ (R1)
Pho92-HTP No UV (R1) Cov.
WT (R1) Cov.
IME2
MEI5
AAGGACACT -160
AAAGACAAT 292
AAGGACACT 89

**Figure S5. (related to Figure 5) Functionality of HTP-tagged proteins used for the CRAC experiment, definition of Pho92 target populations identified by CRAC and enriched motifs, few target examples and correlation with m$^6$A published datasets.**

(**A**) Kinetics of meiotic divisions in different strains of the pCUP-IME1 background used for the CRAC experiment. Meiotic cells correspond to cells containing >1 nucleus as monitored by DAPI staining on 200 cells per time point. Black triangle above 2 hr: time of *IME1* induction.

(**B**) Western blot analysis of different tagged-version of Pho92 across meiosis in WT cells and *ime4Δ/Δ* mutant in the pCUP-IME1 background. HTP- and TAP-tagged version of Pho92 were detected using Peroxidase Anti-Peroxidase complex. Stm1 signal detected in the same extracts was used as loading control. Black triangle above 2 hr: time of *IME1* induction.

(**C**) Western blot analysis of Luc-HTP protein in *S. pombe* extracts grown in rich media. Three increasing quantities of total proteins of the same extracts were used from WT cells, in which no HTP-tag is expressed, and from cells expressing Luc7-HTP, detected with Peroxidase Anti-Peroxidase complex. Oga1, homologous to the *S. cerevisiae* Stm1 protein, was used as loading control and detected with the cross-species reacting anti-Stm1 antibody.

(**D**) Binding of Luc7-HTP to its U1 RNA targets in the CRAC experiment. Left panel: IGV tracks of sequence coverage of the U1 transcript bound by Luc7-HTP in the different samples and replicates (R1, R2) used in the CRAC experiment. Right panel: resulting scores of the U1 peak in the different samples and replicates.

(**E**) Total number of peaks identified with normalized score >0.9 in each sample and biological replicate (R1, R2) used in the CRAC experiment.

(**F**) Pipeline for selection of High-score Ime4-dependent Pho92-binding sites. From the 794 peaks with score >0.9 commonly found in both replicates of Pho92-HTP, was selected those with score >4 in both replicates of Pho92-HTP and a mean score at least 2-fold higher in Pho92-HTP compared to Pho92-HTP in *ime4Δ/Δ*.

(**G**) Distribution of the number of peak(s) per gene for High-score and Lower-score Pho92 targets.

(**H**) Sequence logos of the top ranked motifs identified in the Lower-score Ime4-dependent Pho92 peaks (n= 332), Ime4-independent peaks (n= 142) and unspecific peaks (n= 25), as determined by MEME. The sequences under the peaks of each population was used as input. Only significant motifs are shown with a maximum of 5 motifs and the occurrence of each motif within the population is indicated as well as the associated E-value determined by MEME.

(**I**) Venn diagrams showing the comparison between the genes present in the established methylome datasets in m$^6$A-seq (Schwartz et al 2013), MAZTER-Seq (Garcia-campos et al 2019; group confidence > 1) and Pho92 targets in the High-score peaks or Lower-score peaks. The overlap between genes of the High-score peaks and Lower-score peaks is also shown.

(**J**) Other examples of IGV tracks for Ime4-dependent binding of Pho92-HTP on *IME2* (left) and *MEI5* (right) transcripts. Shown are the sequence coverage (Cov.) for each sample and the crosslink signal (CRAC) detected by the nucleotide deletion induced at the UV-crosslinked binding sites. Only values from one biological replicate are shown for each condition. The identified Pho92-binding peaks are highlighted in grey with the consensus binding sequences indicated at the bottom and the position of the central “A” related to the ATG of each gene.

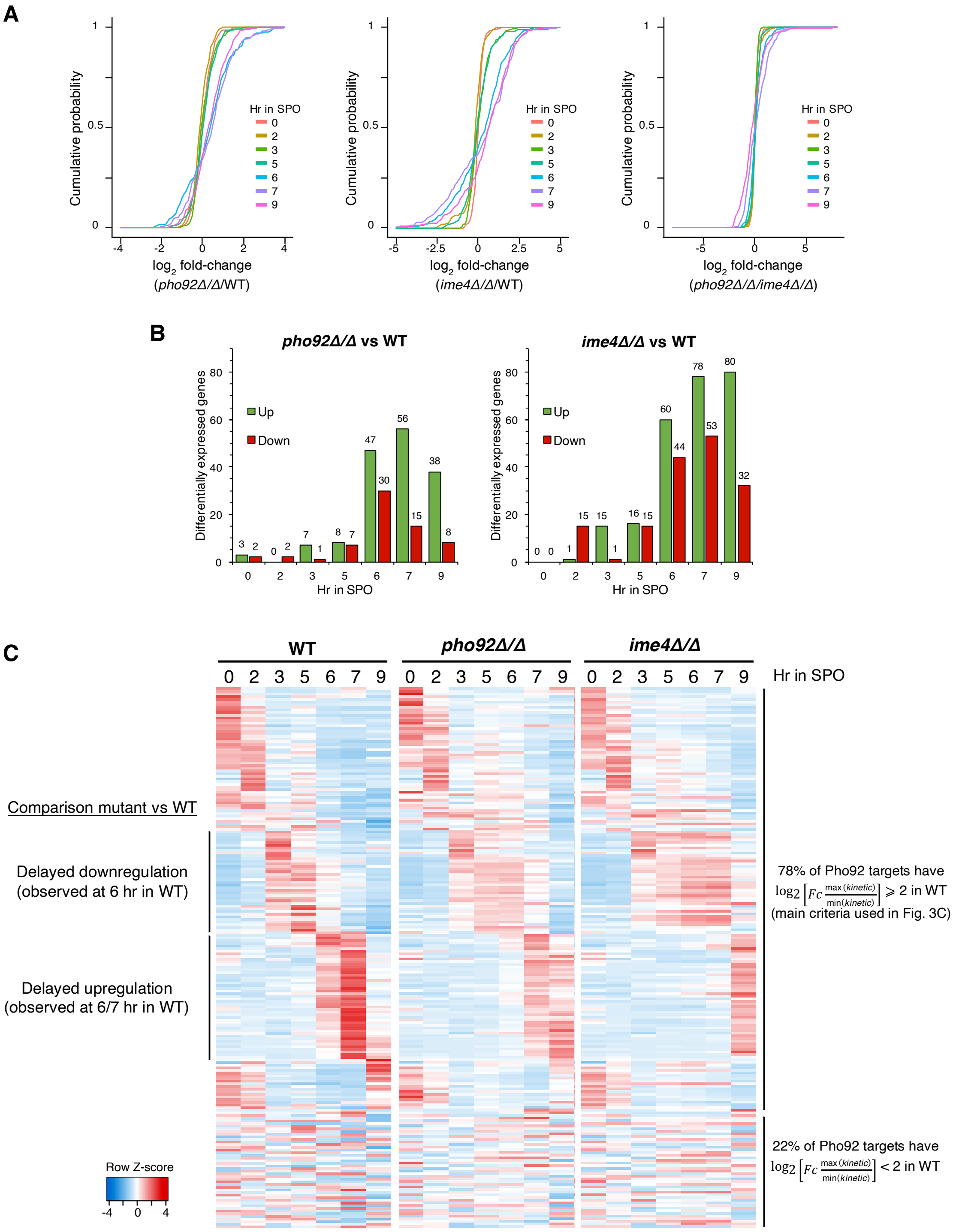


**Figure S6 (related to Figure 6). Expression of the Pho92 targets across meiosis.**

(**A**) Cumulative distribution plots of log2 fold change expression values of the 208 Pho92 High-score targets in *pho92Δ/Δ* over WT (left panel), *ime4Δ/Δ* over WT (middle panel) and *pho92Δ/Δ* over *ime4Δ/Δ* (right panel). Each time point is represented by a different color line.

(**B**) Differential expression of the 208 Pho92 targets in *pho92Δ/Δ* (left panel) and *ime4Δ/Δ* (right panel) mutants compared to WT at each time point of the meiotic time course. Only genes presenting a p-value adjusted for multiple testing <0.05 were considered and defined as up-regulated if log2[Fold change(mutant/WT)] >1 (green) or down-regulated if log2[Fold change(mutant/WT)] <-1 (red). The exact number of genes is indicated on top of each bar.

(**C**) Heat map representation of expression of the 208 High-score Ime4-dependent Pho92 targets during the meiotic time course in WT, *pho92Δ/Δ* and *ime4Δ/Δ* cells. Z-score transformation was performed for each gene considering the mean normalized read count of three biological replicates. Each row represents one gene and each column represents a sample. Z-score is scaled for each line from red (maximum expression value from all samples) to blue (lowest expression value from all samples). Genes are arranged together according to the chronology of their respective maximum expression in the WT strain.
